# Neural Correlates of Motor Imagery, Action Observation, and Movement Execution: A Comparison Across Quantitative Meta-Analyses

**DOI:** 10.1101/198432

**Authors:** Robert M Hardwick, Svenja Caspers, Simon B Eickhoff, Stephan P Swinnen

## Abstract

There is longstanding interest in the relationship between motor imagery, action observation, and movement execution. Several models propose that these tasks recruit the same brain regions in a similar manner; however, there is no quantitative synthesis of the literature that compares their respective networks. Here we summarized data from neuroimaging experiments examining Motor Imagery (303 experiments, 4,902 participants), Action Observation (595 experiments, 11,032 participants), and related control tasks involving Movement Execution (142 experiments, 2,302 participants). Motor Imagery recruited a network of premotor-parietal cortical regions, alongside the thalamus, putamen, and cerebellum. Action Observation involved a cortical premotor-parietal and occipital network, with no consistent subcortical contributions. Movement Execution engaged sensorimotor-premotor areas, and the thalamus, putamen, and cerebellum. Comparisons across these networks highlighted key differences in their recruitment of motor cortex,and parietal cortex, and subcortical structures. Conjunction across all three tasks identified a consistent premotor-parietal and somatosensory network. These data amend previous models of the relationships between motor imagery, action observation, and movement execution, and quantify the relationships between their respective networks.

**Highlights:**

- We compared quantitative meta-analyses of movement imagery, observation, and execution
- Subcortical structures were most commonly associated with imagery and execution
- Conjunctions identified a consistent premotor-parietal-somatosensory network
- These data can inform basic and translational work using imagery and observation

## 1 Introduction

Recent developments have rekindled the longstanding scientific interest in the relationship between the simulation and physical execution of actions. Action simulation (i.e. the internal representation of motor programs without overt movement; Jeannerod, 2001) is typically examined through either motor imagery (i.e. imagining the execution of an action without physically performing it), or action observation (i.e. watching movements performed by others). In particular, motor imagery has received renewed interest following developments in brain computer interface and neurofeedback technology (Liew et al., 2016). This research is supported by decades of work examining the use of motor imagery in elite athletic performance (Calmels et al., 2006; Williams et al., 2015), skill acquisition (Lotze and Halsband, 2006; Pascual-Leone et al., 1995), and rehabilitation (Jackson et al., 2001; but see Ietswaart et al., 2011). Similarly, interest in action observation increased dramatically in the early 2000's following the discovery of 'mirror-neurons' in non-human primates (di Pellegrino et al., 1992; Gallese et al., 1996; Rizzolatti et al., 1996a). Mirror neurons respond both when an action is physically performed, and when the action is observed being performed by another actor. Subsequent research led to considerable investigation of the human action observation system (Grafton et al., 1996; Rizzolatti et al., 1996b). Action observation forms the basis of learning through imitation (Buccino et al., 2004), can induce the same changes in skills as seen in physical practice (Zhang et al., 2011), and is being increasingly examined as a tool for neurorehabilitation (Buccino, 2014; Ertelt et al., 2007). More recent studies have combined mental imagery and action observation (Vogt et al., 2013), allowing greater control over the content and vividness of action simulation (Holmes and Calmels, 2008). Improving our understanding of the brain networks involved in action simulation, and how they relate to the brain regions recruited during movement execution, is therefore of considerable interest to both basic scientific research and translational work across a diverse range of fields.

Several prominent models propose that motor imagery and/or action observation share neural substrates with movement execution (Crammond, 1997; Grèzes and Decety, 2001; Jeannerod, 2001). While early summaries of the literature examined the 'functional equivalence' between motor imagery, action observation, and movement execution, they identified consistent activations across studies in a subjective manner that did not include principled statistical tests (Grèzes and Decety, 2001; Jeannerod, 2001). Later meta-analyses have summarized the individual networks involved in motor imagery (Hétu et al., 2013) and action observation, (Caspers et al., 2010), respectively, but provided no quantitative comparison between their respective networks, or how they compare to the network for movement execution. Questions regarding which regions are consistently implicated in action simulation, and whether there is a consistent network spanning motor imagery, action observation, and movement execution, therefore remain unresolved.

Coordinate-based meta-analysis allows the quantitative summary of the current neuroimaging literature. Pooling data increases statistical power, addressing the limited sample sizes in individual neuroimaging studies. Pooling data also reduces the likelihood of idiosyncratic effects resulting from task-specific activity. Activation Likelihood Estimation (ALE) is an established technique for quantitative voxelwise random effects meta-analysis (Eickhoff et al., 2012, 2009; Laird et al., 2005; Turkeltaub et al., 2012, 2002). Consistently activated regions are determined based on spatial convergence of coordinates reported in previous studies. Statistical testing against a null distribution provides quantitative summary of previous results.

Here we examined the neural correlates of motor imagery, action observation, and movement execution using ALE meta-analysis. We hypothesized that action simulation (i.e. motor imagery and action observation) tasks would recruit broadly similar networks of premotor and parietal brain regions, while movement execution would recruit more sensorimotor regions. Our results identify notable differences and highlight key similarities between these networks, including disparities in their recruitment of subcoritcal structures, and their common recruitment of premotor-parietal and somatosensory regions.

## 2 Methods

### 2.1 *Literature Searches*

Relevant neuroimaging papers were found through pubmed literature searches (as of June 2017). A search for papers on motor imagery was conducted using the search string "((fMRI) OR PET) AND motor imagery", and yielded 487 results. A similar search for papers on action observation was conducted using the search string "((fMRI) OR PET) AND (((action observation) OR mirror neurons) OR imitation)", providing 784 results. The term 'imitation' was included in order to identify contrasts in which participants observed actions prior to imitation. Papers identified in the literature searches were examined for control conditions involving movement execution, allowing us to identify a sample of movement execution tasks with properties similar to those used in the included motor imagery and action observation experiments. This approach reduced the likelihood that differences between the networks were due to inclusion of heterogeneous experimental tasks.

### 2.2 *Inclusion/Exclusion Criteria*

Following the literature searches, inspection of abstracts identified 205 papers on motor imagery and 417 papers on action observation that were downloaded for further inspection. Experiments contained in these papers that used either motor imagery, action observation, or movement execution were assessed for eligibility to be included in the meta-analyses. Only experiments including coordinates from whole brain analyses in standard stereotaxic (MNI/Talairach) space were included in the analyses (to prevent biasing results based on the specific inclusion/exclusion of brain regions). Included experiments reported data from healthy adult participants (i.e. participants ≥18 years of age with no known neurological conditions). Data from healthy control groups in patient studies were included where provided. The meta-analyses examined within-subject contrasts (to prevent comparisons with patient groups, or comparisons across groups of unequal size). Finally, brain activations following neuromodulatory interventions (i.e. measuring the effects of non-invasive brain stimulation or pharmacological agents) were not included, though pre-intervention conditions/control groups were included as appropriate.

### 2.3 *Data Extraction and Classification*

Data extracted from each paper included the number of subjects participating in each experiment, and the coordinates of the reported activations in MNI or Talairach space. Coordinates reported in Talairach space were converted to MNI space using the Lancaster transform (Lancaster et al., 2007). Each task was categorized as involving motor imagery, action observation, and/or movement execution. In order to assess somatotopic activations, we recorded the effector(s) involved in the action, classifying them according to the use of the leg (foot inclusive), arm (hand inclusive), or face (including mouth movements, speech, and facial expressions). Where actions involved multiple effectors they were categorized as using the limbs (both arms and legs, or when contrasts involving the arms and legs were combined), upper body (i.e. movements involving both the face and arm, or contrasts in which face and arm movement were combined), or the whole body (e.g. tasks such as weight lifting or dancing, and conditions in which contrasts involving the leg, arm, and face were combined). Locomotor tasks (including stepping, walking, and running) were categorized as tasks performed with the legs (as the leg acts as the predominant effector). The data included in each meta-analysis and subanalysis are presented in Table 1. More detailed information on the individual experiments included in each meta-analysis is presented in Supplementary Table 1.

**Table 1:** Data included in the meta-analyses

| **Analysis** | **Experiments** | **Participants** | **Foci** |
| --- | --- | --- | --- |
| Motor Imagery | 303 | 4902 | 3235 |
| *Somatotopy subanalyses:* |  |  |  |
| - Leg | 65 | 916 | 801 |
| - Arm | 179 | 3041 | 1928 |
| - Face* | 6 | 111 | 57 |
| Action Observation | 595 | 11032 | 6561 |
| *Somatotopy Subanalyses:* |  |  |  |
| - Leg | 34 | 453 | 297 |
| - Arm | 339 | 6494 | 3831 |
| - Face | 64 | 1103 | 761 |
| Movement Execution | 142 | 2302 | 1842 |
| *Somatotopy Subanalyses:* |  |  |  |
| - Leg | 20 | 208 | 239 |
| - Arm | 107 | 1858 | 1324 |
| - Face* | 13 | 219 | 214 |

\* Analysis should be considered exploratory as it includes less than 20 experiments (Eickhoff et al., 2016)

### 2.4 *Data Analyses*

In a first step we conducted ALE meta-analyses to identify the individual task networks involved in motor imagery, action observation, and movement execution. Each task network was examined in greater detail using subanalyses including experiments using only the face, only the arm, and only the foot as an effector. We then assessed the convergence and divergence between the individual task networks. Networks were overlaid in a pairwise fashion, allowing us to identify the volume of each network that was specific to each individual task, and the volume that was activated across multiple tasks. We identified regions that were consistently engaged across different tasks by computing pairwise conjunction analyses, and in a final step a combined conjunction identified the regions consistently recruited across all three tasks.

### 2.5 *Analysis Procedure*

All analyses were conducted using the revised version of the activation likelihood estimation (ALE) algorithm (Eickhoff et al., 2009; Turkeltaub et al., 2002). The ALE approach empirically determines whether converging activation coordinates (foci) across different experiments occurs at a level greater than expected by chance. Reported foci are modeled as the centers of 3D Gaussian probability distributions (Turkeltaub et al., 2002). The revised algorithm sets the width of these Gaussians using empirical between-subject and between-template comparisons, and models the increased spatial reliability of larger sample sizes by using smaller Gaussian distributions (Eickhoff et al., 2009). Comparisons between groups of different sizes are accounted for by computing a null-distribution using label-exchangeability (Eickhoff et al., 2012).

Foci for each experiment were combined across voxels to produce a modeled activation map (Turkeltaub et al., 2012). Combining modeled activation maps across experiments produced ALE scores, which described the convergence of coordinates for each location. ALE scores were compared to a non-linear histogram integration based on the frequency of distinct modeled activation maps (Eickhoff et al., 2012), determining areas where convergence is greater than expected by chance. ALE values were computed only for voxels with a ≥10% probability of containing grey matter (Evans et al., 1994), as functional activations occur predominantly in grey matter areas. Results were thresholded at p<0.05 (cluster-level FWE, corrected for multiple comparisons, cluster-forming threshold at voxel level p<0.001) and provided at 2mm^3^ voxel resolution.

Contrasts between the resultant meta-analyses were conducted using random effects ALE subtraction analysis (Eickhoff et al., 2012). In a first step, voxel-wise differences between ALE maps were calculated for each pool of experiments. Experiments were then randomly shuffled into two samples of equal size to the compared analyses, and voxelwise differences between their ALE scores were recorded. This shuffling procedure was repeated 10,000 times to produce an empirical null distribution of ALE score differences between the compared conditions. The map of differences based on this procedure was thresholded at a posterior probability for true differences of P>0.95, and inclusively masked by the respective main effect of the minuend (cf. Chase et al., 2011; Rottschy et al., 2012) with a minimum cluster volume of 100mm^3^ (Beissner et al., 2013; Erickson et al., 2014; Turkeltaub et al., 2012).

Volume comparisons were conducted by overlaying the networks and determining the number of voxels that were unique to each analysis, or co-recruited across analyses. As differences in the number of studies included in the meta-analyses could influence the size of the volume identified in each case, we also conducted 'volume matched analyses'. Using a previously established procedure we iteratively increased the threshold for the larger network, reducing its size until the number of voxels it identified was approximately equivalent to that of the smaller network (Hardwick et al., 2015). This identified those regions most consistently implicated in a paradigm while controlling for differences in the number of studies included in each analysis.

Conjunction analyses were conducted to summarize the overlap between networks. These analyses used the conjunction null hypothesis and were calculated using the minimum statistic (Nichols et al., 2005), with a minimum cluster volume of 100mm^3^ (Beissner et al., 2013; Erickson et al., 2014; Turkeltaub et al., 2012). In a final step we conducted a conjunction across the analyses of motor imagery, action observation, and movement execution to identify neural substrates commonly recruited by all three tasks.

### 2.6 *Labeling*

Results were anatomically labeled according to their most probable macroanatomical and cytoarchitectonic/tractographically assessed locations using the SPM Anatomy Toolbox 2 extension (Eickhoff et al., 2007, 2006, 2005). Additional functional labels for motor and premotor cortical regions were identified using the human motor area template (HMAT) as defined by Mayka et al. (2006). Coordinates were reported based on peak maxima in MNI space.

## 3 Results

### 3.1 *Meta Analyses*

In a first step we conducted quantitative meta-analysis of motor imagery, movement execution, and movement execution (Figure 1).

**Figure 1:**
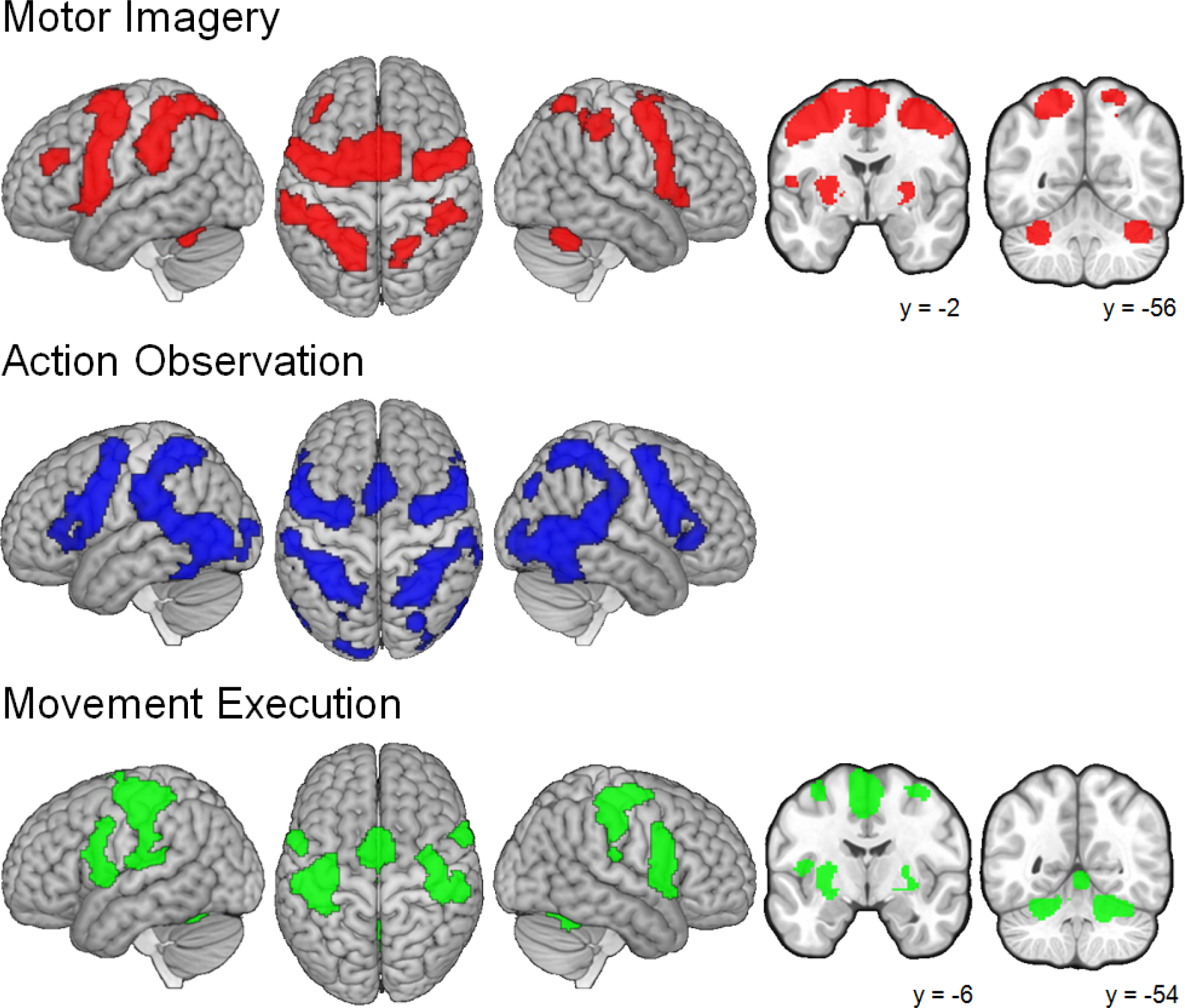
*Quantitative meta-analyses of the three tasks. Note that no slices are shown for action observation as the meta-analysis did not identify clusters with maxima in subcortical areas*.

#### 3.1.1 *Motor Imagery*

**Table 2:**
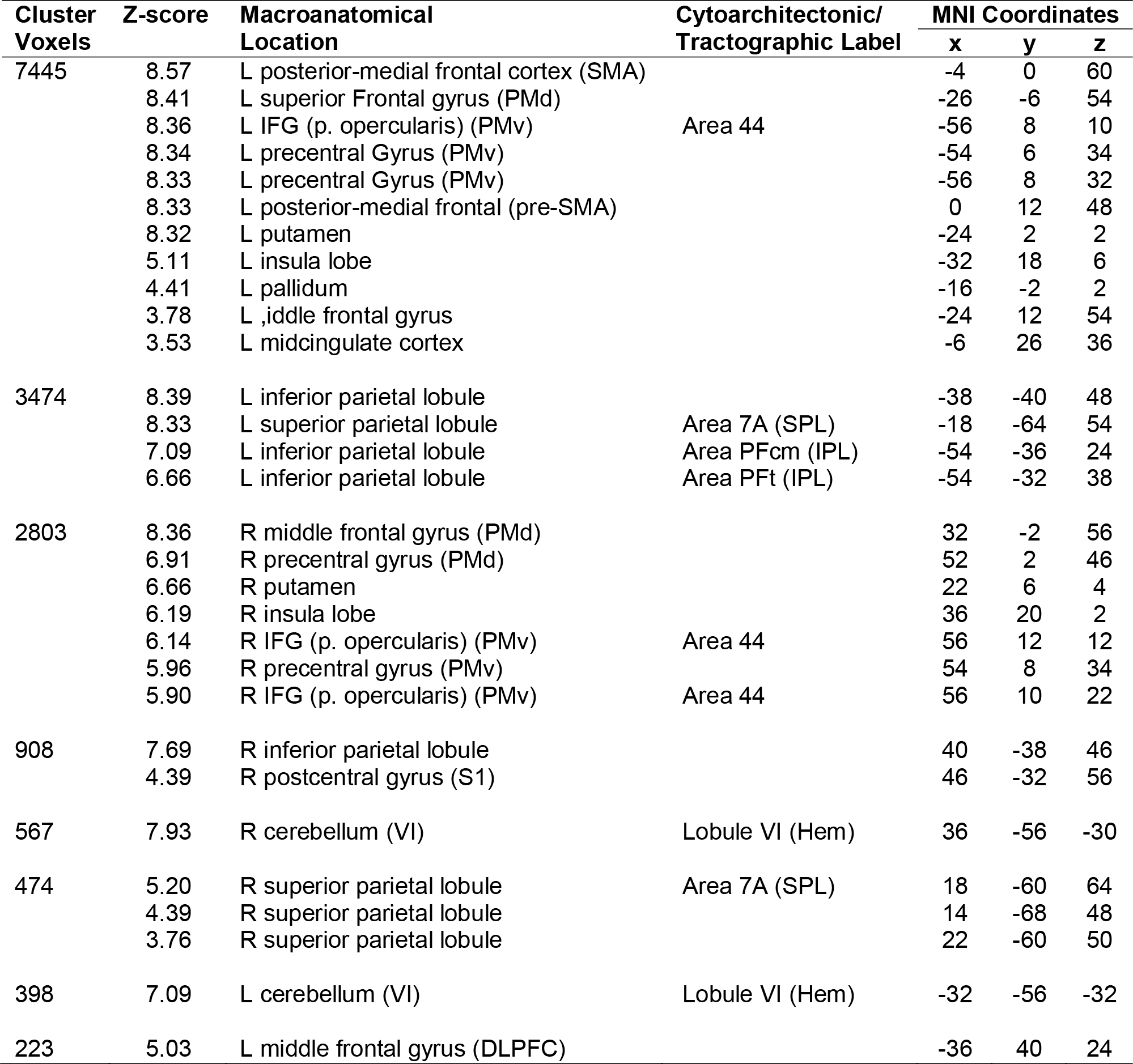
Peak coordinates identified by the meta-analysis of motor imagery

Motor Imagery primarily recruited a network of bilateral premotor, rostral inferior and middle superior parietal, basal ganglia, and cerebellar regions. Two large bilateral premotor clusters spanned the SMA-proper and pre-SMA, extending to the dorsal and ventral premotor cortices. The left premotor cluster also extended to encompass areas of the cingulate and putamen (notably, a separate smaller cluster also included the right putamen). Two bilateral parietal clusters spanned the inferior and superior parietal lobules, with right lateralized activations in the inferior parietal sulcus. Further subcortical clusters were identified biltaterally in lobule VI of the cerebellum. There was also a relatively small cluster in the left middle frontal gyrus consistent with the dorsolateral prefrontal cortex (Al-Hakim et al., 2006).

#### 3.1.2 *Action Observation*

**Table 3:** Peak coordinates from the meta-analysis of action observation

| Cluster Voxels | Z-score | Macroanatomical Location | Cytoarchitectonic/ Tractographic Label | MNI Coordinates |  |  |
| --- | --- | --- | --- | --- | --- | --- |
|  |  |  |  | x | y | z |
| 7381 | 8.86 | L middle occipital gyrus | hOc5 [V5/MT] | -48 | -72 | 2 |
|  | 8.60 | L superior parietal lobule | Area 7PC (SPL) | -34 | -46 | 56 |
|  | 8.42 | L inferior parietal lobule | Area PFt (IPL) | -56 | -26 | 36 |
|  | 8.42 | L inferior temporal gyrus |  | -42 | -46 | -18 |
|  | 8.42 | L inferior parietal lobule | Area PFcm (IPL) | -52 | -38 | 24 |
|  | 8.42 | L middle temporal gyrus |  | -52 | -48 | 8 |
|  | 5.70 | L superior occipital gyrus | hOC1 [V1] | -10 | -100 | 6 |
|  | 5.61 | L inferior occipital gyrus | hOc3v [V3v] | -22 | -90 | -10 |
|  | 4.62 | L middle occipital gyrus | hOc3d [V3d] | -24 | -96 | 10 |
|  | 4.13 | L superior occipital gyrus |  | -24 | -74 | 40 |
| 7126 | 8.88 | R middle temporal gyrus | hOc5 [V5/MT] | 48 | -66 | 0 |
|  | 8.51 | R inferior parietal lobule |  | 34 | -42 | 54 |
|  | 8.48 | R middle temporal gyrus |  | 54 | -42 | 8 |
|  | 8.46 | R superior temporal gyrus |  | 60 | -36 | 18 |
|  | 8.42 | R fusiform gyrus |  | 42 | -48 | -18 |
|  | 5.68 | R inferior occipital gyrus |  | 38 | -84 | -4 |
|  | 5.49 | R inferior parietal lobule |  | 58 | -22 | 40 |
|  | 5.07 | R middle occipital gyrus | hOc3v [V3v] | 28 | -94 | 0 |
|  | 4.80 | R lingual gyrus | hOc3v [V3v] | 26 | -90 | -8 |
|  | 4.25 | R fusiform gyrus | hOc4v [V4(v)] | 28 | -78 | -10 |
| 3575 | 8.68 | L superior frontal gyrus (PMd) |  | -26 | -6 | 54 |
|  | 8.50 | L IFG (p. opercularis) (PMv) | Area 44 | -48 | 10 | 24 |
|  | 8.47 | L precentral gyrus (PMv) |  | -52 | 6 | 36 |
|  | 8.43 | L precentral gyrus (PMd) |  | -48 | -2 | 46 |
|  | 7.42 | L insula lobe |  | -34 | 20 | 0 |
|  | 4.78 | L IFG (p. triangularis) | Area 45 | -50 | 32 | 10 |
|  | 4.71 | L IFG (p. triangularis) | Area 44 | -52 | 16 | 4 |
|  | 3.61 | L IFG (p. orbitalis) |  | -44 | 28 | -4 |
|  | 3.47 | L IFG (p. triangularis) |  | -52 | 30 | -2 |
| 2959 | 8.50 | R precentral gyrus (PMv) |  | 52 | 8 | 32 |
|  | 8.48 | R precentral gyrus (PMd) |  | 30 | -6 | 56 |
|  | 8.43 | R precentral gyrus (PMd) |  | 46 | 2 | 48 |
|  | 5.76 | R insula lobe |  | 34 | 22 | -2 |
|  | 5.12 | R IFG (p. triangularis) | Area 45 | 54 | 30 | 0 |
|  | 3.91 | R IFG (p. triangularis) | Area 45 | 48 | 30 | 16 |
|  | 3.71 | R IFG (p. orbitalis) |  | 46 | 24 | -4 |
| 893 | 8.39 | L posterior-medial frontal (pre-SMA) |  | -2 | 12 | 52 |
|  | 5.14 | L posterior-medial frontal |  | -2 | 6 | 62 |
| 210 | 6.89 | R superior occipital gyrus |  | 26 | -80 | 36 |
|  | 3.42 | R middle occipital gyrus |  | 30 | -78 | 26 |

The action observation meta-analysis identified the network with the greatest overall volume of the three meta-analyses conducted. Similarly to the imagery network, action observation recruited a bilateral network of premotor and parietal regions; however, this network included large parietal-occipital volumes, and identified more bilateral clusters. Two large bilateral clusters spanned the dorsal and ventral premotor cortices, while a third smaller premotor cluster was identified in the bilateral pre-SMA. A smaller cluster included the right superior occipital gyrus.The two largest clusters identified by this analysis covered bilateral parietal-occipital regions, spanning from the superior parietal lobule down to the inferior parietal lobule, and portions of the occipital cortex. Notably, these parieto-occipital clusters included some minor bilateral excursions into the bilateral cerebellum that did not include peak maxima. It is thus notable that action observation did not lead to any consistent recruitment of subcortical regions.

#### 3.1.3 *Movement execution*

**Table 4:** Local maxima from the meta-analysis of movement execution

| Cluster Voxels | Z-score | Macroanatomical Location | Cytoarchitectonic/ Tractographic Label | MNI Coordinates |  |  |
| --- | --- | --- | --- | --- | --- | --- |
|  |  |  |  | x | y | z |
| 3110 | 8.38 | L precentral gyrus (M1) | Area 4a | -38 | -22 | 56 |
|  | 7.10 | L postcentral gyrus (S1) | Area 2 | -50 | -24 | 46 |
|  | 6.43 | L rolandic operculum | Area OP 1 | -46 | -24 | 18 |
|  | 5.77 | L inferior parietal lobule |  | -50 | -26 | 28 |
|  | 5.16 | L inferior parietal lobule |  | -60 | -22 | 16 |
|  | 3.53 | L inferior parietal lobule | Area PFt (IPL) | -60 | -32 | 32 |
| 1775 | 8.39 | L posterior-medial frontal (SMA) |  | -2 | 0 | 54 |
|  | 4.72 | L midcingulate cortex |  | -4 | 8 | 40 |
|  | 4.68 | L posterior-medial frontal cortex (SMA) |  | -6 | -22 | 50 |
| 1668 | 8.31 | L rolandic operculum (PMv) |  | -48 | 2 | 6 |
|  | 7.59 | L putamen |  | -26 | -6 | 0 |
|  | 7.15 | L precentral gyrus (PMv) | Area 44 | -56 | 8 | 28 |
|  | 4.35 | L precentral gyrus (PMv) |  | -54 | 0 | 40 |
|  | 3.84 | L IFG (p. opercularis) |  | -50 | 16 | 20 |
| 1441 | 8.31 | R cerebellum (VI) | Lobule VI (Hem) | 20 | -54 | -24 |
|  | 6.30 | Cerebellar vermis (VI) | Lobule VI (Hem) | 6 | -64 | -18 |
|  | 5.78 | Cerebellar vermis (IV/V) | Lobule V (Hem) | 2 | -56 | -4 |
|  | 5.23 | R cerebelum (IV-V) | Lobule V (Hem) | 16 | -40 | -24 |
| 1347 | 6.50 | R precentral gyrus (M1) |  | 40 | -18 | 60 |
|  | 5.62 | R postcentral gyrus (S1) |  | 42 | -26 | 60 |
|  | 4.90 | R postcentral Gyrus (S1) | Area 1 | 54 | -20 | 44 |
|  | 4.55 | R rolandic operculum |  | 50 | -24 | 20 |
|  | 4.32 | R inferior parietal lobule |  | 58 | -30 | 44 |
|  | 4.26 | R inferior parietal lobule |  | 52 | -24 | 34 |
| 1115 | 8.04 | R insula |  | 48 | 6 | 4 |
|  | 6.45 | R IFG (p. opercularis) | Area 44 | 58 | 10 | 28 |
|  | 5.93 | R IFG (p. opercularis) | Area 44 | 60 | 12 | 12 |
|  | 4.95 | R temporal pole |  | 54 | 14 | -6 |
| 615 | 7.52 | L cerebelum (VI) | Lobule VI (Hem) | -24 | -54 | -24 |
|  | 3.16 | L cerebelum (IV-V) |  | -6 | -52 | -18 |
| 383 | 8.33 | L thalamus | Thal: Prefrontal | -14 | -20 | 6 |
| 249 | 5.23 | R putamen |  | 24 | -4 | 8 |
|  | 4.42 | R pallidum |  | 22 | -4 | -6 |
|  | 4.35 | R pallidum |  | 24 | -2 | -4 |
|  | 4.19 | R pallidum |  | 26 | -4 | -2 |
| 193 | 7.46 | R thalamus | Thal: Premotor | 14 | -18 | 6 |

The network identified by the movement execution meta-analysis was the smallest in volume of the three meta-analyses. Cortical activations spanned the sensorimotor and premotor cortices, and included small regions of the inferior parietal lobule, while subcortical clusters were identified in the bilateral thalamus, putamen, and cerebellum. Though larger in the left hemisphere, two bilateral clusters spanned the primary motor and primary somatosensory cortex, with anterior regions reaching into the dorsal premotor cortex. Premotor convergence was identified in three clusters; a cluster spanning the bilateral SMA (extending down to the cingulate cortex), and two bilateral clusters across the ventral premotor cortex. Consistent with the bilateral but primarily left-lateralized clusters in the sensorimotor cortex, subcortical activity included bilateral (primarily left lateralized) clusters in the thalamus, and bilateral (primarily right-lateralized) clusters in cerebellar lobule VI.

### 3.2 *Subanalyses*

A series of subanalyses were conducted on experiments in which the tasks were performed predominantly with the leg, arm, or face (Figure 2). These analyses aimed to better characterize the volumes for each task, and to probe for somatotopic organization within the identified networks.

**Figure 2:**
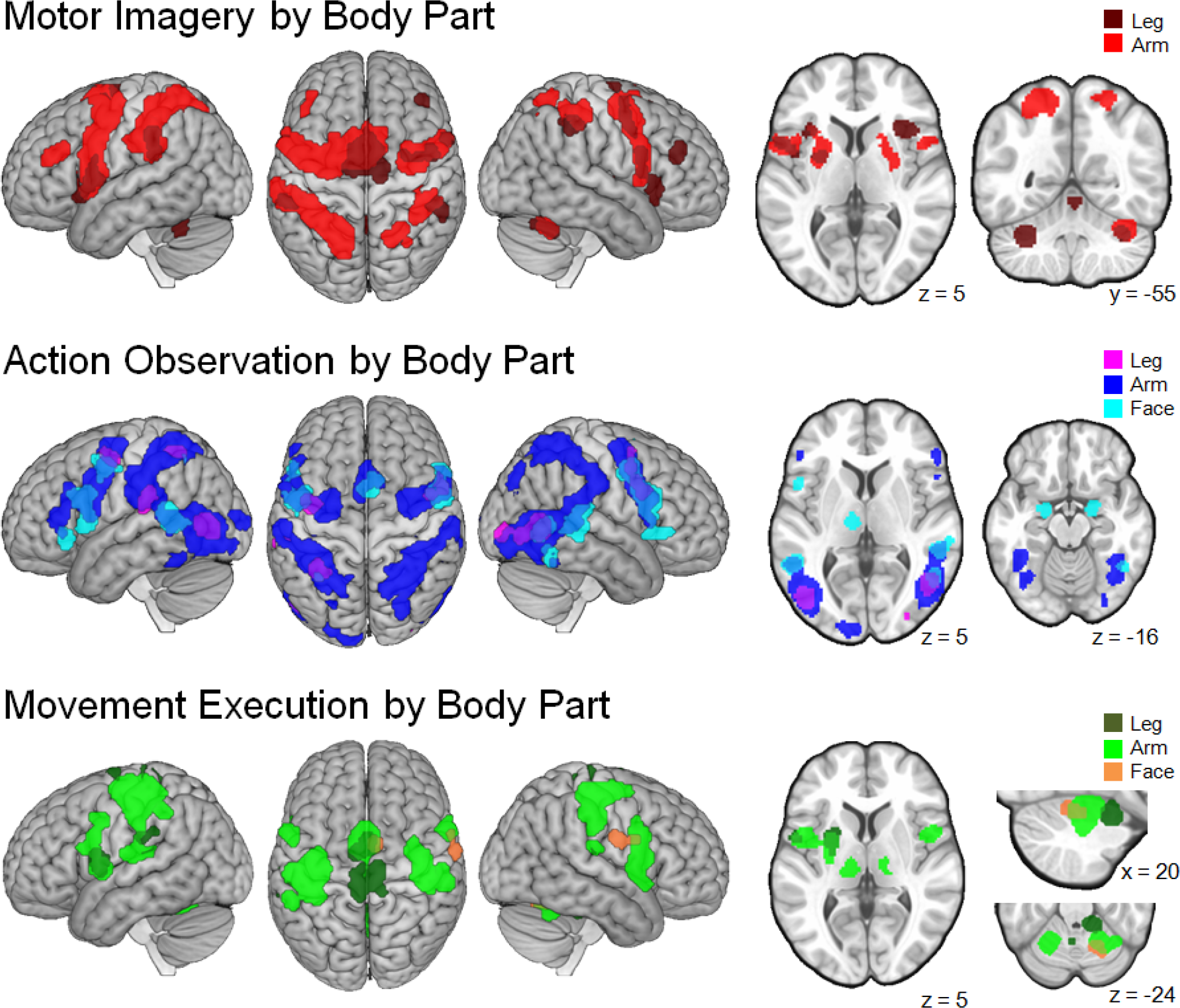
Subanalyses for each task conducted according to the body part used.

#### 3.2.1 *Motor Imagery by Body Part*

Results for the subanalyses examining only imagery tasks performed with either the leg or arm were broadly consistent with those of the main analysis of motor imagery. The exploratory analysis of the six studies examining motor imagery with the face identified no converging activations. Notably, tasks performed with the leg recruited the right dorsolateral prefrontal cortex. Motor imagery with both the leg and arm recruited a similarly located volume in the right cerebellum; tasks performed with the leg also recruited clusters in the left cerebellum and cerebellar vermis, most likely due to the bilateral nature of the included tasks. While previous studies have reported that motor imagery recruits premotor and parietal regions in a somotatopic manner (Stippich et al., 2002), there was little evidence of such organization within the present results.

#### 3.2.2 *Action Observation by Body Part*

Subanalyses of action observation tasks performed using the leg, arm, and face identified sub-networks consistent with the main analysis. This analysis identified limited evidence of somatotopic organization in the premotor and parietal lobes, with a greater likelihood of superior regions being recruited by observing the leg, and a greater likelihood for inferior regions being recruited when observing the face. The subanalyses also identified a region of the right extrastriate visual cortex responsive to actions performed with the leg, arm, or face. Notably, contrary to the main analysis, the subanalysis of action observation tasks involving the face identified consistent recruitment of the left thalamus and bilateral amygdala.

#### 3.2.3 *Movement Execution by Body Part*

The subanalyses performed on movement execution tasks performed with the leg and arm, and the exploratory subanalysis of the 12 studies involving movements of the face, identified a network similar to that found in the main analysis. An interesting feature of these subanalyses was their somatotopic recruitment of cortical and subcortical areas. In the primary motor cortex, activity along the central sulcus was consistent with classic motor mappings (Penfield and Rasmussen, 1950), with a superior cluster associated with leg activity, a relatively central cluster associated with arm activity (including the region of M1 associated with the hand; Yousry et al., 1997), and a more inferior cluster associated with the face. The right cerebellum was also recruited in a somatotopic manner; tasks performed with the leg, arm, or face corresponded to relatively anterior, central, and posterior regions, respectively (Figure 2, lower panel).

### 3.3 *Contrast Analyses*

ALE contrast analyses identified regions more consistently implicated with one of the tasks when compared to another (Figure 3).

**Figure 3:**
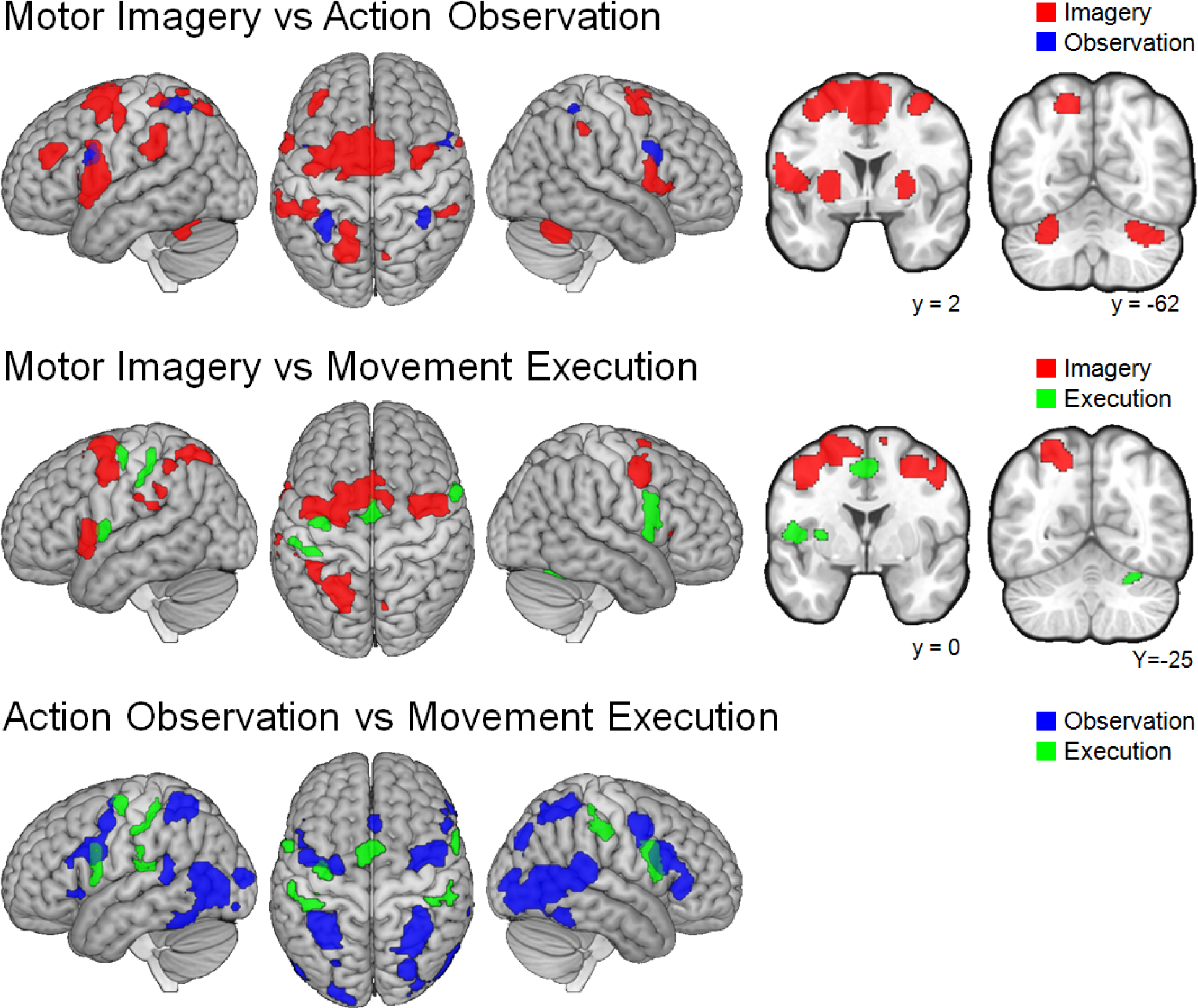
*Contrast analyses. Note that small regions of apparent 'overlap' present areas that are rendered on the cortical surface, but are present at different depths. No slices are shown for the comparison between action observation and movement execution as no subcortical clusters were identified in the analysis*.

#### 3.3.1 *Motor Imagery vs Action Observation*

Motor imagery was more consistently associated with recruiting a majority of premotor regions, including bilateral SMA, PMd, and PMv. Imagery was also more consistently linked with parietal regions, recruiting bilateral areas of the inferior parietal lobe, and regions of the (mainly left) superior parietal lobe. The left DPMFC was also more likely to be recruited by imagery than action observation. At a subcortical level the bilateral putamen and cerebellum were both more consistently linked with motor imagery tasks in comparison to action observation.

Relatively few clusters were identified as being more likely to be recruited during action observation when compared to motor imagery. These areas included relatively small, bilateral regions of the inferior frontal gyrus and areas of the right inferior/superior parietal lobule (Notably, these regions correspond to the premotor and parietal regions in which mirror-neurons have been identified in non-human primates - see Fogassi et al., 2005; Gallese et al., 1996).

#### 3.3.2 *Motor Imagery vs Movement Execution*

Motor imagery was, when compared to movement execution, more consistently associated with recruiting premotor regions, including the bilateral pre-SMA, PMd, and left PMv. A large area of the left parietal cortex, spanning the inferior and superior parietal lobule, was also more consistently associated with motor imagery than movement execution. No subcortical regions were more closely associated with motor imagery than movement execution.

By contrast, movement execution was, in comparison to motor imagery, more consistently implicated with classic sensorimotor regions including the SMA-proper and cingulate motor areas, left primary motor and somatosensory cortex, and bialteral ventral premotor cortex. At a subcortical level movement execution was associated with recruitment of the left putamen and lobule VI of the right cerebellum.

#### 3.3.3 *Action Observation vs Movement Execution*

Action observation, in comparison to movement execution, recruited a mainly bilateral network of premotor, parietal, and occipital regions. This included the Pre-SMA, bilateral PMd and PMv, bilateral inferior and superior parietal lobe, and bilateral visual cortex. No subcortical regions were more closely associated with action observation when compared to movement execution.

Regions more consistently associated with movement execution than action observation included a mainly bilateral cortical sensorimotor network. This included the SMA proper, left primary motor cortex, bilateral somatosensory cortex, and bilateral ventral premotor cortex. No subcortical regions were found to be more consistently implicated in movement execution than action observation.

### 3.4 *Volume Comparisons*

In a series of volume comparisons, we quantified the extent to which each identified volume was unique, and to which it overlapped with other analyses (Figure 4). This analysis therefore quantified the similarity and disparity between each identified network.

**Figure 4:**
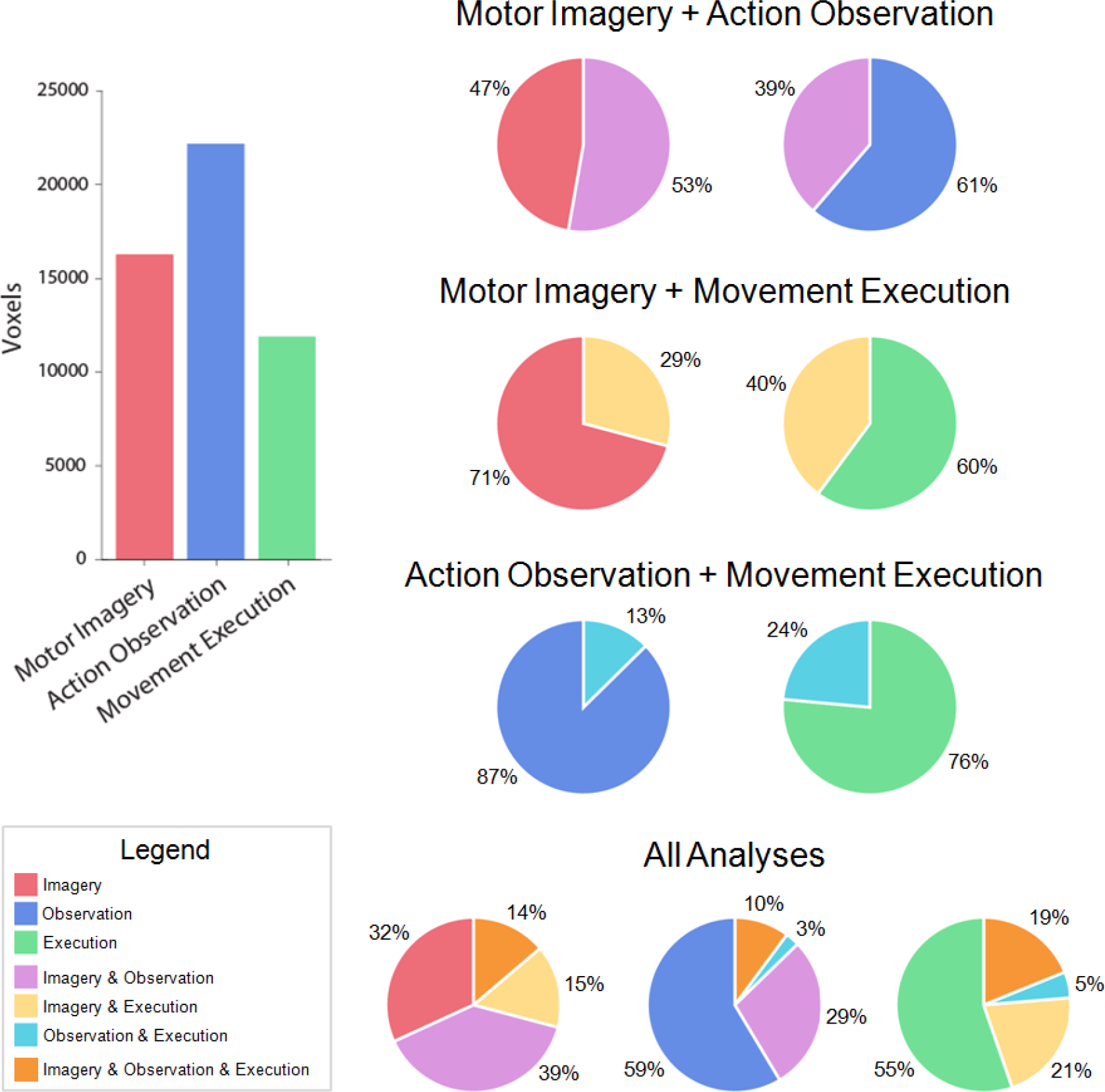
*Volume Comparisons. Bar chart illustrates the number of voxels contributing to the volume for each task. Pie charts illustrate the percentage of each volume that is unique to one task, or overlaps with other tasks. Note that the bar chart illustrates that the overall volume of each network differs; thus, the same absolute volume of overlap represents a different percentage of the corresponding individual networks*.

#### 3.4.1 *Imagery and Observation*

There was considerable overlap between the volumes involved in action simulation. The majority (53%) of voxels identified as being involved in motor imagery were also identified as being involved in action observation. Motor imagery also recruited 39% of the volume identified by action observation.

As the volume of the motor imagery network was smaller than that for action observation, we conducted a secondary volume matched analysis. We iteratively increased the threshold of the larger (observation) analysis until the volume of voxels approximately matched that of the imagery analysis. This resulted in a 43% overlap between each network.

#### 3.4.2 *Imagery and Execution*

The networks involved in motor imagery and movement execution were also relatively comparable. 29% of voxels identified in the analysis of motor imagery were also identified as being involved in movement execution. Imagery also recruited 40% of the voxels that were identified in the movement execution analysis. Matching these volumes to the smaller (movement execution) result indicated that 33% of voxels for each analysis were involved in both modalities.

#### 3.4.3 *Observation and Execution*

The overlap between observation and execution (i.e. the analysis for potential 'mirror' areas) was smaller than those identified in the analyses above. For the volume involved in observation, only 13% of voxels were also involved in execution. In comparison, observation recruited 24% of voxels identified by the movement execution analysis. As the volume of the observation network was almost twice that of the execution network, more strict thresholding of the observation analysis such that it matched the execution analysis indentified that only 14% of the resulting volumes were active across both analyses.

#### 3.4.4 *Imagery, Observation, and Execution*

The majority of the volume identified by motor imagery was also recruited by at least one other task. While 32% of the volume was uniquely recruited during motor imagery, a larger proportion (39%) was recruited by motor imagery and action observation. Smaller proportions of the imagery volume were also active during movement execution (15%), or were active across all three tasks (14%).

The volume identified by action observation was generally unimodal; 59% of the voxels were recruited by action observation alone. There was greater overlap between action observation and motor imagery (29% than action observation and movement execution (3%). In total, 10% of the voxels involved in action observation was recruited by all three tasks.

Most of the voxels recruited during movement execution were also unimodal, with 55% of the volume responding only during movement execution. There was a greater overlap between movement execution and motor imagery (21%) than movement execution and action observation (5%), though 19% of the volume recruited by movement execution was involved in all three tasks.

These results changed little when the volume of the motor imagery and action observation networks were matched to the smaller volume of the movement execution analysis. For the motor imagery volume, 39% was unique to imagery, 29% overlapped with observation alone, 21% overlapped with execution, and 12% overlapped across all three tasks. For action observation, 57% of the volume was unimodal, 29% overlapped only with motor imagery, 3% overlapped only with movement execution, and 12% overlapped across all three tasks. Finally, for movement execution, 65% of the volume was not active in other tasks, 21% was activated only during both motor imagery and movement execution, 3% overlapped between action observation and movement execution, and 12% overlapped across all tasks.

### 3.5 *Conjunction Analyses*

Models comparing motor imagery, action observation, and movement execution have proposed that they all recruit a shared network of brain regions. similar networks of brain regions. Here we tested this hypothesis by conducting a series of conjunction analyses using the minimum statistic to identify regions consistently recruited by more than one task (Figure 5).

**Figure 5:**
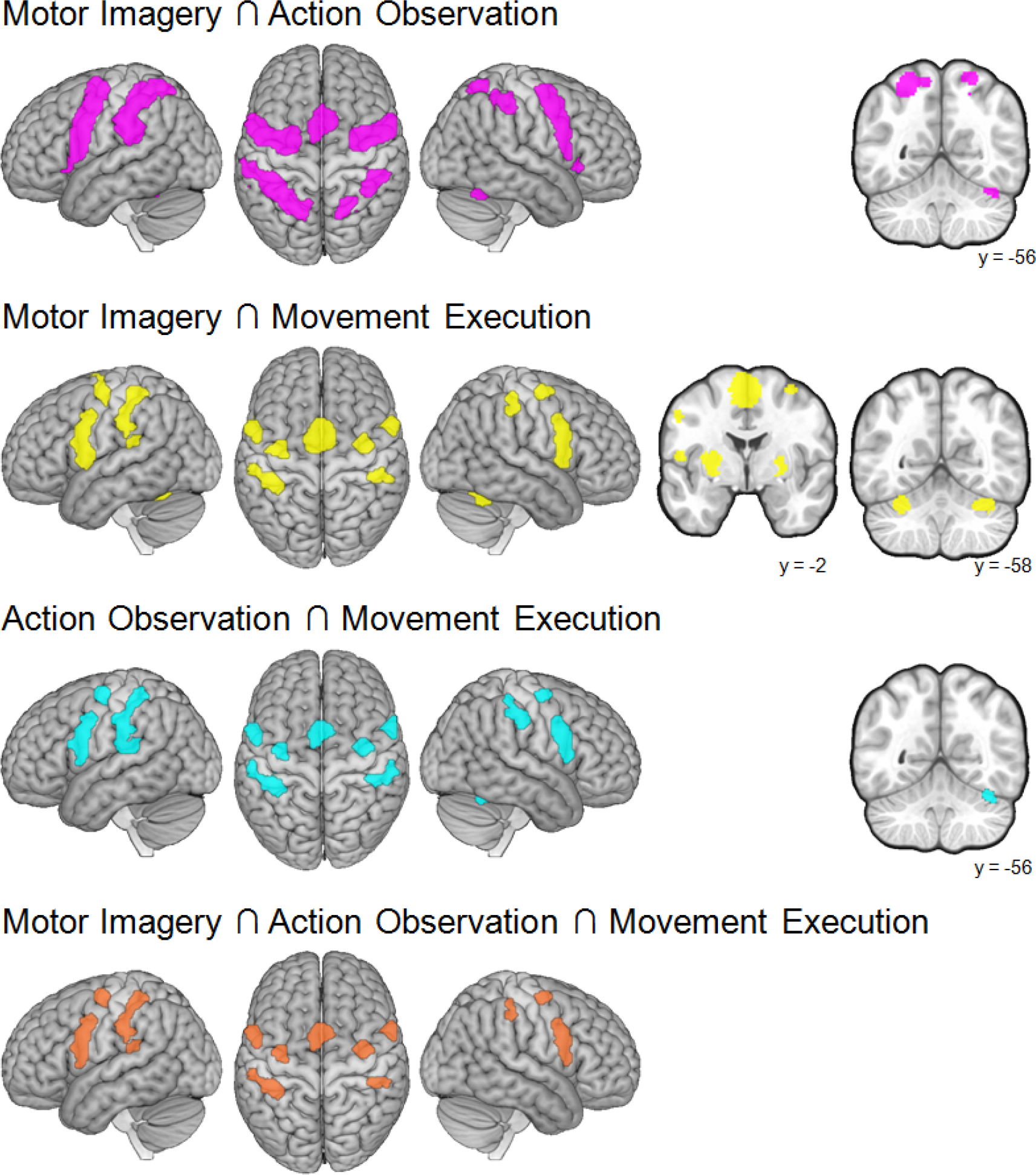
Conjunction analyses conducted across combinations of the tasks. Note that slices are only shown in cases where analyses identified subcortical clusters.

#### 3.5.1 *Motor Imagery* ∩ *Action Observation*

**Table 5:** Peak coordinates from the conjunction between motor imagery and action observation

| Cluster Voxels | Z-score | Macroanatomical Location | Cytoarchitectonic/<br>Tractographic Label | MNI Coordinates |  |  |
| --- | --- | --- | --- | --- | --- | --- |
|  |  |  |  | x | y | Z |
| 2512 | 8.39 | L inferior parietal lobule |  | -38 | -40 | 48 |
|  | 7.09 | L inferior parietal lobule | Area PFcm (IPL) | -54 | -36 | 24 |
|  | 6.48 | L inferior parietal lobule | Area PFt (IPL) | -54 | -30 | 38 |
|  | 6.15 | L superior parietal lobule | Area 7A (SPL) | -22 | -60 | 58 |
|  | 5.95 | L superior parietal lobule | Area 7A (SPL) | -24 | -54 | 64 |
| 2304 | 8.41 | L superior frontal gyrus (PMd) |  | -26 | -6 | 54 |
|  | 8.34 | L precentral gyrus (PMv) |  | -54 | 6 | 34 |
|  | 8.32 | L precentral gyrus (PMv) | Area 44 | -56 | 8 | 28 |
|  | 8.04 | L IFG (p. opercularis) (PMv) | Area 44 | -54 | 10 | 14 |
|  | 7.29 | L precentral gyrus (PMd) |  | -42 | -2 | 52 |
| 1584 | 8.36 | R middle frontal gyrus (PMd) |  | 32 | -2 | 56 |
|  | 6.91 | R precentral gyrus (PMd) |  | 52 | 2 | 46 |
|  | 5.96 | R precentral gyrus (PMv) |  | 54 | 8 | 34 |
|  | 5.82 | R IFG (p. opercularis) (PMv) | Area 44 | 54 | 10 | 22 |
|  | 5.24 | R IFG (p. opercularis) (PMv) | Area 44 | 52 | 12 | 12 |
| 805 | 7.99 | L posterior-medial frontal (pre-SMA) |  | -2 | 10 | 52 |
|  | 5.14 | L posterior-medial frontal (pre-SMA) |  | -2 | 6 | 62 |
| 703 | 7.69 | R intraparietal sulcus | Area hIP2 (IPS) | 40 | -38 | 46 |
| 280 | 4.96 | R superior parietal lobule | Area 7A (SPL) | 18 | -62 | 62 |
|  | 3.75 | R superior parietal lobule |  | 14 | -66 | 52 |
|  | 3.66 | R intraparietal sulcus | Area hIP3 (IPS) | 20 | -60 | 52 |
|  | 3.59 | R superior parietal lobule |  | 24 | -60 | 50 |
| 188 | 4.84 | L insula |  | -32 | 20 | 4 |
|  | 3.81 | L insula |  | -40 | 16 | 2 |
| 119 | 5.13 | R insula |  | 34 | 20 | 0 |
| 98 | 4.98 | R cerebellum (VI) |  | 40 | -56 | -24 |

A conjunction between motor imagery and action observation identified areas consistently recruited during action simulation. This action simulation network included primarily bilateral premotor and rostral parietal regions, with greater cortical volumes being identified in the left hemisphere. A premotor cluster spanning the midline was identified within the pre-SMA, and further bilateral clusters spanned the dorsal and ventral premotor cortex for each hemisphere. Bilateral clusters were identified across the superior and inferior parietal lobules. Notably, a relatively small cluster was as having a peak maximum in the right cerebellar hemisphere. However, further inspection identified that the region from the action observation meta analysis that contributed to this cluster primarily corresponded to the extra-striate visual area, and featured a small incursion into the cerebellum without peak maxim. This indicates that the cerebellar cluster identified in the conjunction analysis may not have been driven by shared Cerebellar recruitment per-se, but could instead be attributed to some coincidental overlap between adjacent clusters.

#### 3.5.2 *Motor Imagery* ∩ *Movement execution*

**Table 6:**
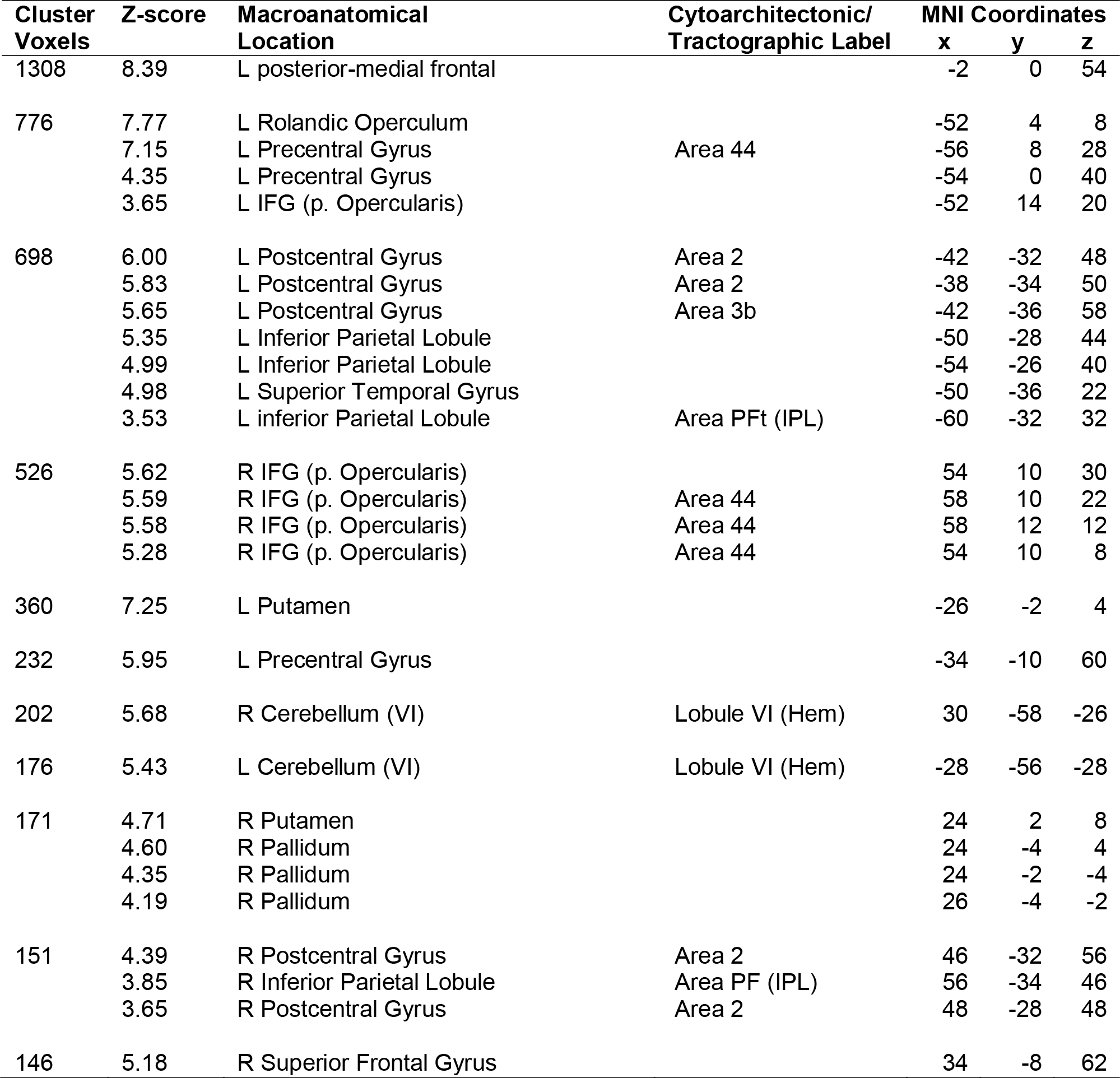
Local maxima from the conjunction between motor imagery and movement execution

A conjunction between motor imagery and movement execution identified a network including bilateral cortical sensorimotor and premotor clusters, with subcortical clusters of lesser volume in the putamen and cerebellum. In premotor regions, one cluster included the bilateral pre-SMA and SMA proper, and a further small cluster was identified in the right dorsal premotor cortex, while further bilateral clusters included the ventral premotor cortex. Posterior to these premotor clusters were two bilateral clusters that each included the primary somatosensory cortex. Subcortically, the bilateral putamen and cerebellum (lobule VI) were also identified as consistent across both constituent analyses.

### 3.6 *Action Observation* ∩ *Movement Execution*

**Table 7:** Results of the conjunction between action observation and movement execution

| Cluster Voxels | Z-score | Macroanatomical Location | Cytoarchitectonic/ Tractographic Label | MNI Coordinates |  |  |
| --- | --- | --- | --- | --- | --- | --- |
|  |  |  |  | x | y | z |
| 957 | 7.05 | L Postcentral Gyrus | Area 3b | -38 | -32 | 50 |
|  | 5.96 | L inferior Parietal Lobule |  | -54 | -24 | 42 |
|  | 5.65 | L Postcentral Gyrus | Area 3b | -42 | -36 | 58 |
|  | 5.41 | L Superior Temporal Gyrus |  | -50 | -34 | 18 |
|  | 5.32 | L inferior Parietal Lobule | Area PFt (IPL) | -52 | -26 | 30 |
|  | 5.26 | L inferior Parietal Lobule | Area PFop (IPL) | -50 | -28 | 28 |
|  | 5.00 | L inferior Parietal Lobule |  | -48 | -30 | 26 |
|  | 4.07 | L inferior Parietal Lobule |  | -58 | -24 | 18 |
|  | 3.53 | L inferior Parietal Lobule | Area PFt (IPL) | -60 | -32 | 32 |
| 480 | 7.15 | L Precentral Gyrus | Area 44 | -56 | 8 | 28 |
|  | 6.17 | L IFG (p. Opercularis) | Area 44 | -54 | 6 | 12 |
|  | 4.35 | L Precentral Gyrus |  | -54 | 0 | 40 |
|  | 3.85 | L IFG (p. Opercularis) |  | -50 | 16 | 20 |
| 440 | 6.57 | L posterior-medial frontal |  | -2 | 8 | 54 |
| 343 | 6.45 | R IFG (p. Opercularis) | Area 44 | 58 | 10 | 28 |
|  | 4.65 | R IFG (p. Opercularis) | Area 44 | 58 | 16 | 10 |
| 229 | 4.49 | R Postcentral Gyrus |  | 56 | -22 | 42 |
|  | 4.30 | R Postcentral Gyrus | Area 2 | 44 | -34 | 56 |
|  | 4.15 | R Postcentral Gyrus | Area 3b | 40 | -32 | 56 |
|  | 4.01 | R inferior Parietal Lobule | Area PFt (IPL) | 52 | -30 | 46 |
| 169 | 5.79 | L Precentral Gyrus |  | -34 | -12 | 56 |
| 117 | 4.85 | R Superior Frontal Gyrus |  | 36 | -8 | 60 |
| 60 | 4.39 | R Cerebellum (VI) | Lobule VI (Hem) | 38 | -54 | -24 |

Consistent activations across action observation and movement execution were identified in a bilateral premotor, parietal, and sensorimotor and premotor network. Premotor regions included the bilateral pre-SMA extending back into the left SMA-proper, separate clusters in the bilateral ventral premotor cortex, and a small cluster in the right dorsal premotor cortex. Parietal convergence spanned the inferior parietal lobule. A small cluster was identified in the right cerebellum; however, as in the conjunction of motor imagery and action observation, the further inspection of the action observation meta-analysis indicated that the contributing cluster originated in the visual cortex. Again, this is consistent with the cluster identified in the conjunction analysis not being a result of direct cerebellar recruitment in both constituent analyses; instead this may reflect coincidental overlap between adjacent clusters.

### 3.7 *Motor Imagery* ∩ *Action Observation* ∩ *Movement Execution*

**Table 8:** Results of the conjunction across all three tasks (motor imagery, action observation, and movement execution).

| Cluster Voxels | Z-score | Macroanatomical Location | Cytoarchitectonic/<br>Tractography Label | MNI Coordinates |  |  |
| --- | --- | --- | --- | --- | --- | --- |
|  |  |  |  | x | y | z |
| 633 | 6.00 | L Postcentral Gyrus | Area 2 | -42 | -32 | 48 |
|  | 5.83 | L Postcentral Gyrus | Area 2 | -38 | -34 | 50 |
|  | 5.65 | L Postcentral Gyrus | Area 3b | -42 | -36 | 58 |
|  | 5.23 | L Inferior Parietal Lobule |  | -52 | -28 | 44 |
|  | 4.99 | L Inferior Parietal Lobule |  | -54 | -26 | 40 |
|  | 4.98 | L Superior Temporal Gyrus |  | -50 | -36 | 22 |
|  | 3.53 | L inferior Parietal Lobule | Area PFt (IPL) | -60 | -32 | 32 |
| 459 | 7.15 | L Precentral Gyrus | Area 44 | -56 | 8 | 28 |
|  | 6.17 | L IFG (p. Opercularis) | Area 44 | -54 | 6 | 12 |
|  | 4.35 | L Precentral Gyrus |  | -54 | 0 | 40 |
|  | 3.65 | L IFG (p. Opercularis) |  | -52 | 14 | 20 |
| 440 | 6.57 | L posterior-medial frontal |  | -2 | 8 | 54 |
| 298 | 5.62 | R IFG (p. Opercularis) |  | 54 | 10 | 30 |
|  | 4.50 | R IFG (p. Opercularis) | Area 44 | 56 | 14 | 10 |
| 162 | 5.67 | L Precentral Gyrus |  | -34 | -10 | 58 |
| 116 | 4.85 | R Superior Frontal Gyrus |  | 36 | -8 | 60 |
| 89 | 4.02 | R Postcentral Gyrus | Area 2 | 44 | -34 | 56 |
|  | 3.85 | R Postcentral Gyrus | Area PFt (IPL) | 50 | -30 | 46 |

A grand conjunction across motor imagery, action observation, and movement execution identified brain areas involved in both the simulation and performance of actions. This analysis identified a bilateral network of premotor, parietal, and sensory regions. Separate premotor clusters spanned the pre-SMA and left SMA-proper, and the bilateral dorsal and ventral premotor cortices. More posterior clusters included bilateral regions of parietal and sensorimotor cortex.

## 4 Discussion

### 4.1 *Task networks*

Here for the first time we quantified and compared the extent of the individual networks involved in motor imagery, action observation, and movement execution. This allowed us to assess the networks involved in action simulation tasks, and determine how they relate to the network involved in movement execution. As previous meta-analyses have examined the individual networks for motor imagery (Hétu et al., 2013) and action observation (Caspers et al., 2010), our discussion focuses on notable differences and similarities between these networks, and on their relation to the network for movement execution.

#### 4.1.1 *Motor Imagery*

The meta-analysis of motor imagery identified a predominantly premotor-parietal network, with subcortical recruitment of the putamen and cerebellum. While our volume comparison analysis identified the greatest overlap between motor imagery and action observation, this was mainly limited to premotor and parietal regions. Later conjunction analyses identified a more diverse network of regions were both active during motor imagery and movement execution, including the midcingulate cortex, putamen, and cerebellum.

Notably, the motor imagery meta-analysis was unique in identifying consistent recruitment of the dorsolateral prefrontal cortex (DLPFC), and corresponding regions of the frontal thalamus. This is in accord with the previously established role of the DLPFC in working memory, with a particular emphasis on spatial working memory (Barbey et al., 2013). The DLPFC is also implicated in frontal-executive functions related to action preparation (Mars and Grol, 2007), which is believed to have similar neural substrates to motor imagery. However, the DLPFC was not recruited during action observation or movement execution. DLPFC recruitment during motor imagery could therefore be mostly related to increased demands on working memory, as evidence indicates this can be separate to frontal-executive DLPFC functions recruited during more complex movement tasks (Rottschy et al., 2012; Wollenweber et al., 2014). Alternatively, as the DLPFC plays a role in movement inhibition (Blasi et al., 2006; Nigel et al., 2015), it may act to inhibit overt movement execution during motor imagery.

Both motor imagery and movement execution recruited areas of the midcingulate cortex. Motor imagery recruited a relatively anterior region proposed to play an important role in the more cognitive aspects of motor control (Hoffstaedter et al., 2013a). By contrast, movement execution recruited a relatively posterior region more directly associated with motor function (Picard and Strick, 1996; Procyk et al., 2014) Conjunction across the analyses identified a large cluster with peak maxima in the SMA that extended down into to the midcingulate cortex, but had no peak maxima within the cingulate itself. The majority of the overlap was in relatively posterior cingulate regions, consistent with a role in movement production.

Motor imagery and movement execution both recruited bilateral areas of the putamen. A region of the basal ganglia that forms a critical node of the cortico-striatal sensorimotor circuit (Voon et al., 2015), the putamen is associated with automatic movement behaviors (Ashby and Crossley, 2012). Activity in the putamen correlates with the speed and extent of executed movements (Turner et al., 2003), consistent with evidence that the basal ganglia are involved in calculating the cost of movements (Shadmehr and Krakauer, 2008). This is notable as disorders of the basal ganglia related to Parkinson's disease lead patients to become slower to move (Dickson, 2017), and lead to increases in the duration of imagined movements (Helmich et al., 2007; Heremans et al., 2011). The putamen may therefore be involved in regulating the speed of self paced imagined or executed actions. This could also explain why the basal ganglia were not implicated in action observation; the viewer has no ability to regulate the speed of the observed action.

The cerebellum was recruited during both motor imagery and movement execution; in particular, the right cerebellar lobule VI was consistently involved in both tasks as identified by a conjunction analysis. The highly consistent nature of the right cerebellar cluster is in accordance with previous observations of cerebellar-thalamo-motor connectivity (Buckner et al., 2011; Daskalakis et al., 2004). The cerebellum contains multiple representations of the body, and results from the meta-analysis of movement execution showed somatotopic effects in line with previous work (Buckner et al., 2011; Debaere et al., 2001). Cerebellar lobule VI contains body representations that are most prominent during movement execution (Schlerf et al., 2010). However, our subanalysis of motor imagery identified little evidence of somatotopic recruitment of the cerebellum. This may suggest that the function for which the cerebellum was recruited differed between motor imagery and movement execution (discussed further below).

#### 4.1.2 *Action Observation*

The action observation meta-analysis identified a cortical network of mainly premotor-parietal and occipital regions. Occipital regions were uniquely associated with action observation, which was unsurprising given the use of visual stimuli in the included contrasts. Notably, a subanalysis examining action observation studies according to the limb being observed identified three notable features of the network. First, we identified a right hemisphere extrastriate visual region that responded to the presentation of actions performed across the leg, arm, or face. This is consistent with previously described sensitivity of this extrastriate region to the observation of movement and parts of the body (Born and Bradley, 2005; Downing et al., 2001; Ferri et al., 2013; Urgesi et al., 2007); for a review see Lingnau and Downing, 2015). Second, the subanalysis identified bilateral recruitment of the amygdala in response to observing faces, consistent with the emotional content of the facial stimuli presented in many of the included experiments (Carr et al., 2003; Grosbras and Paus, 2005; Lenzi et al., 2009; van der Gaag et al., 2007). This is in line with work indicating that action observation can include empathic components (Avenanti et al., 2005). Finally, this subanalysis also provided limited evidence of somatotopic organization within the premotor and parietal cortex, in line with previous work (Buccino et al., 2001; Jastorff et al., 2010; Lorey et al., 2013).

In a contrast analysis, motor imagery was more consistently associated with recruiting a wide range of premotor and parietal regions than action observation. There were, however, notable exceptions to this, with small clusters in the inferior frontal gyrus (ventral premotor cortex) and inferior/superior parietal cortex being more closely associated with action observation than motor imagery. This is notable as the regions identified are highly consistent with the areas in which mirror neurons have been identified in nonhuman primates (di Pellegrino et al., 1992; Fogassi et al., 2005; Rizzolatti et al., 1996a).

A notable result was that the main analysis of action observation did not identify consistent recruitment of subcortical areas. While a cluster spanning parietal and extrastriate visual areas did include a relatively small area of the cerebellum, no peak maxima were identified within the cerebellum itself. This result is consistent with a previous meta-analysis of action observation that found no evidence of consistent recruitment of the cerebellum (Caspers et al., 2010).

#### 4.1.3 *Movement Execution*

Movement execution recruited a cortical sensorimotor and premotor network, with further subcortical clusters in the putamen, thalamus, and cerebellum. While this analysis included relatively few studies, it was highly consistent with several previously identified hallmarks of the motor system. Specifically, subanalyses identified somatotopic recruitment of the primary motor cortex and cerebellum, consistent with established motor maps within these structures (Buckner et al., 2011; Penfield and Rasmussen, 1950; Schlerf et al., 2010). The finding that this network also recruited the thalamus is consistent with the well established role of this structure as a cortico-subcortical relay (Sommer, 2003). Finally, we note that the network for movement execution identified here recruited a similar volume to that identified in our previous meta-analysis of motor learning (Hardwick et al., 2013). Notably, only the movement execution network was found to consistently recruit the primary motor cortex (see below for further discussion).

### 4.2 *Consistent sub-network for Motor Imagery, Action Observation, and Movement Execution*

A conjunction across motor imagery, action observation, and movement execution identified a network of premotor, rostral parietal, and somatosensory regions.

#### 4.2.1 *Premotor Cortex*

The bilateral ventral premotor cortex, dorsal premotor cortex, and pre-SMA were all consistently implicated in motor imagery, action observation, and movement execution. These regions of premotor cortex are typically associated with action preparation (Hoshi and Tanji, 2007), and with linking arbitrary stimuli or conditions to actions (Nachev et al., 2008; Wise And Murray, 2000). The ventral premotor cortex interacts with the primary motor cortex to shape the hand during grasping actions (Davare et al., 2009; Rizzolatti et al., 1988), suggesting PMv plays an important role in fine motor coordination. Work in primates indicates that the macaque homologue of PMv contains 'mirror neurons', sensitive to both the observation and execution of actions (di Pellegrino et al., 1992; Gallese et al., 1996; Rizzolatti et al., 1996a). Similar to PMv, PMd has reciprocal connections with M1 and the spinal cord, but has limited ability to directly contribute to movement execution (Boudrias et al., 2010; Dum and Strick, 2005). PMd is therefore believed to play more of a role in action selection than execution (Halsband et al., 1993; Rushworth et al., 1998). A particularly notable property of PMd neurons is that their firing patterns change as primates learn arbitrary associations between stimuli and actions (Wise and Murray, 2000). The visuomotor associative properties of premotor cortex have been examined in studies that have established 'mirror' and 'counter-mirror' responses in ventral and dorsal premotor areas through training (Catmur et al., 2011, 2008). Similarly, SMA is associated with linking conditional rules to actions (Nachev et al., 2008), and is important to self-initiated actions (Deecke and Kornhuber, 1978; Hoffstaedter et al., 2013b). The more anterior pre-SMA regions identified in the grand conjunction analysis are consistently associated with both motor tasks, and with non-motor cognitive processes (Leek and Johnston, 2009; Tanaka et al., 2005), consistent with the demands required during motor imagery tasks. The multimodal properties of these premotor regions are therefore consistent with their activation across motor imagery, action observation, and motor execution tasks.

#### 4.2.2 *Parietal Cortex*

The bilateral inferior parietal lobule region PFt was consistently activated across all analyses. The parietal cortex is typically involved in processing multisensory information (Block et al., 2013), and PFt is involved in the performance of motor behaviors and the processing of tactile information (Klann et al., 2015), with a special role in tool use (Orban, 2016). Human area PFt has been proposed as the homologue of primate area PF (Caspers et al., 2010), which contains mirror neurons (Fogassi et al., 2005). The consistent co-activity of the parietal and premotor cortices identified by the conjunction analyses is consistent with their interactions during visuomotor control of actions (Wise et al., 1997). Patients with damage to the parietal lobule also have impairments in using mental imagery to accurately predict the time required to perform motor tasks (Sirigu et al., 1996).

While large portions of the parietal cortex were involved in the networks for motor imagery and action observation, it was involved to a much lesser extent in movement execution. Notably, the movement execution tasks in the present sample were relatively simple, and evidence from lesion studies indicates that parietal damage has more pronounced effects on complex actions (De Renzi et al., 1983; Weiss et al., 2008). Therefore, the relative simplicity of the actions performed in the movement execution conditions may contribute to this effect.

#### 4.2.3 *Somatosensory Cortex*

The recruitment of somatosensory regions during motor imagery is consistent with the kinesthetic aspects of motor imagery (i.e. imagining the sensations associated with performing actions). Models of action observation have also proposed that 'mirror' properties extend beyond the motor system to those involved in movement execution (Keysers and Gazzola, 2009). The prominent 'threshold theory' of mirror-touch synesthesia (a condition in which observing another person being touched evokes the sensation of being touched) proposes that somatosensory mirror activity occurs in all individuals, only becoming mirror-touch synesthesia once it passes the threshold for active perception (Ward and Banissy, 2015). In movement execution, sensory information provides critical feedback for the accuracy of movements, allowing comparison of the actual and predicted sensory consequences of actions (Hardwick et al., 2012; Muckli and Petro, 2017). This finding is also consistent with models proposing that motor imagery, and potentially action observation, lead to sensory efference in a similar manner as movement execution (Crammond, 1997).

### 4.3 *Comparisons with previous models*

Jeannerod's (2001) simulation theory proposed the same network of brain regions are recruited during motor imagery, action observation, and movement execution. The proposed simulation network included the primary motor cortex, corticospinal pathway, basal ganglia, cerebellum, premotor cortex, parietal cortex, and prefrontal cortex. The majority of these areas were found to be active across at least two of the tasks identified in our analyses; however, we found only truly consistent recruitment of the premotor and rostral parietal cortices across all tasks, and in addition identified converging activity in the primary somatosensory cortex. The premotor and parietal network identified here is therefore broadly consistent with previous models proposing that motor imagery, action observation, and movement execution recruit the same neural structures (Crammond, 1997; Grèzes and Decety, 2001; Jeannerod, 2001).

The involvement of the primary motor cortex in action simulation has long been a subject of debate. Jeannerod (2001) argued that fMRI results show clear involvement of M1 in action simulation. In contrast, Grèzes and Decety (2001) proposed that the recruitment of M1 during action simulation was ambiguous; they noted that PET studies generally showed no involvement of M1, while fMRI studies did report recruitment of sensorimotor cortex. However, meta-analyses of neuroimaging data provide evidence against the recruitment of M1 during action simulation. Caspers et al., (2010) found evidence that M1 may only be recruited during action observation when participants view actions with the intention to imitate them. Similarly, Hétu et al., (2013) found no evidence of consistent recruitment of M1 during motor imagery. The results of the present study are consistent with the view that M1 is only recruited during movement execution. However, we note early reviews reported that increases in M1 activity reported during action simulation tasks were less than those seen during movement execution (Grèzes and Decety, 2001; Jeannerod, 2001), and that it has been proposed that M1 may be active at a level lower than to induce peak maxima (Lotze and Halsband, 2006).

Studies using TMS to show increases in corticospinal excitability during motor imagery and action observation have also been cited as evidence that M1 is recruited during action simulation (for a review see Loporto et al., 2011). Similarly, later studies of action observation in non-human primates have provided evidence of mirror neurons in M1 (Dushanova and Donoghue, 2010; Tkach et al., 2007; Vigneswaran et al., 2013). However, these data provide conflicting results; while early TMS studies indicated increases in M1 excitability, recordings from the primate corticospinal tract indicate that neurons in M1 show a suppression of their firing rates in response to observing actions (Vigneswaran et al., 2013). This discrepancy indicates that these TMS results are unlikely to result from direct increases in M1 excitability, but could result from the premotor cortex acting to increase MEP amplitudes through cortico-cortical or cortico-subcortical pathways (Fadiga et al., 2005).

Conjunctions across the networks identified relatively little consistent recruitment of subcortical structures. Action observation did not consistently recruit the subcortical structures, and, contrary to proposed models, did not appear to directly recruit the cerebellum (Miall, 2003). Notably, motor imagery and movement execution did both recruit regions of the thalamus, putamen, and cerebellum. This runs counter to earlier reports that proposed the basal ganglia were not recruited during motor imagery (for review see Jackson et al., 2001). This difference may be due to earlier studies not providing whole-brain coverage due to a-priori hypotheses and technical limitations. The consistent recruitment of subcortical structures associated with movement execution during motor imagery, but not during action observation, may have important implications for translational research (see below).

#### 4.3.1 *Is Co-activation Across Tasks Evidence of Functional Equivalence?*

Previous studies have proposed that overlapping activations during motor imagery, action observation, and movement execution provide evidence of 'functional equivalence' within these regions; that the same regions perform the same computations across the three different tasks. This stems from an intuitive assumption that it would be inefficient to develop distinct networks for these similar tasks (Hétu et al., 2013), and could explain the majority of the overlap between the networks identified in our analyses.

It is, however, difficult to infer the exact role a region plays in a task on the basis of neuroimaging data (Poldrack, 2006). Furthermore, activation across multiple tasks is not necessarily consistent with functional equivalence. For example, our subanalysis of motor imagery indicated that the same region of the cerebellum was recruited regardless of whether the task was performed with the leg, arm, or foot, whereas our subanalysis of movement execution identified clear somatotopic recruitment of the cerebellum. Thus, while a conjunction across tasks identified a region of the cerebellum that was consistently implicated in both motor imagery and action observation, it does not necessarily indicate that the cerebellum functions in the same manner across both tasks. Furthermore, the traditional BOLD contrast used in fMRI studies, and the haemodynamic response as measured in PET studies, provide only indirect evidence of functional equivalence. As each voxel contains many thousands of neurons, it is possible that co-activation across tasks could be due to the responses of separate sub-populations. This limitation is partially addressed through the use of fMRI repetition suppression - this approach assumes that the firing response of neurons will attenuate if they are consistently presented with a stimulus to which they are sensitive. A number of studies attempting to identify mirror neurons in the human brain have probed for 'cross-modal' repetition suppression between action observation and movement execution with mixed results (for a review see Kilner and Lemon, 2013). Unfortunately, many of these studies used a limited search volume restricted to premotor regions, and were therefore not suitable to be included in the present meta-analyses. We therefore conclude that the co-activation across motor imagery, action observation, and movement execution observed in the present analyses is consistent with the theory of functional equivalence. However, further research using approaches more sensitive to the activity of individual neurons, and/or multivariate approaches that allow more sensitive assessments of the action representations involved in motor imagery, action observation, and movement execution (e.g. Zabicki et al., 2017) is still required to confirm this hypothesis.

#### 4.3.2 *Implications for translational research*

Results of the present meta-analyses indicate that while motor imagery and action observation recruit a consistent network of premotor and parietal areas, motor imagery recruited more regions that were also involved in movement execution. This suggests that translational research with the aim of recruiting circuits involved in movement execution may be best supported by interventions using motor imagery, as action observation recruited less of the motor network. However, for clinical settings, there is evidence that imagined actions can be limited by the same impairments that affect movement execution (Helmich et al., 2007; Heremans et al., 2011). Such limitations could be overcome by combining motor imagery and action observation. This combined approach could engage circuits implicated in movement execution, while concurrently allowing accurate control of the content of imagined actions (Holmes and Calmels, 2008). Such a combination could therefore be of particular interest in clinical settings (Eaves et al., 2014; Vogt et al., 2013).

While there is considerable interest in using action simulation in rehabilitation settings, it has thus far provided mixed results. Small scale studies in stroke provide promising preliminary data (Ertelt et al., 2007), however, larger clinical trials indicate no effects (Ietswaart et al., 2011). Considering that even extensive physical training has limited impact on motor abilities in the chronic phase of recovery (Hardwick et al., 2017; Kitago and Krakauer, 2013), it would be expected that interventions using action simulation would have minimal effects. Notably, the majority of recovery in stroke occurs not during the chronic phase, but in the acute period within the first three months after stroke (Xu et al., 2017). Work in animal models suggests physical training early after stroke may improve overall recovery (Zeiler et al., 2015; Zeiler and Krakauer, 2013). As patients are not cleared for physical activity in the early acute phase, action simulation may therefore provide an alternate approach to promote early motor recovery.

## 5 Conclusions

Previous comparisons between motor imagery, action observation, and movement execution have relied on the results of individual neuroimaging studies, or qualitative comparisons of their respective networks. Here for the first time we provide a quantitative comparison of the brain networks involved in these similar tasks. Our results provide an empirical answer to the longstanding debate over whether these tasks recruit similar brain regions by identifying a consistent premotor, parietal, and somatosensory network.

## Acknowledgements

We thank Stefan Vogt (Lancaster) for insightful comments on an earlier draft of this manuscript. We also thank Sebastien Hétu (Virginia Tech) and Philip Jackson (Laval) for sharing data from their meta-analysis of motor imagery.

## References

Al-Hakim, R., Fallon, J., Nain, D., Melonakos, J., Tannenbaum, A., 2006. A dorsolateral prefrontal cortex semi-automatic segmenter, in: Reinhardt, J.M., Pluim, J.P.W. (Eds.),. International Society for Optics and Photonics, p. 61440J. doi:10.1117/12.653643

Ashby, F.G., Crossley, M.J., 2012. Automaticity and multiple memory systems. Wiley Interdiscip. Rev. Cogn. Sci. 3, 363–376. doi:10.1002/wcs.1172

Avenanti, A., Bueti, D., Galati, G., Aglioti, S.M., 2005. Transcranial magnetic stimulation highlights the sensorimotor side of empathy for pain. Nat. Neurosci. 8, 955–960. doi:10.1038/nn1481

Barbey, A.K., Koenigs, M., Grafman, J., 2013. Dorsolateral prefrontal contributions to human working memory. doi:10.1016/j.cortex.2012.05.022

Beissner, F., Meissner, K., Bär, K.-J., Napadow, V., 2013. The Autonomic Brain: An Activation Likelihood Estimation Meta-Analysis for Central Processing of Autonomic Function. J. Neurosci. 33, 10503–11. doi:10.1523/JNEUROSCI.1103-13.2013

Blasi, G., Goldberg, T.E., Weickert, T., Das, S., Kohn, P., Zoltick, B., Bertolino, A., Callicott, J.H., Weinberger, D.R., Mattay, V.S., 2006. Brain regions underlying response inhibition and interference monitoring and suppression. Eur. J. Neurosci. 23, 1658–1664. doi:10.1111/j.1460-9568.2006.04680.x

Block, H., Bastian, A., Celnik, P., 2013. Virtual lesion of angular gyrus disrupts the relationship between visuoproprioceptive weighting and realignment. J. Cogn. Neurosci. 25, 636–48. doi:10.1162/jocn_a_00340

Born, R.T., Bradley, D.C., 2005. Structure and function of visual area MT. Annu. Rev. Neurosci. 28, 157–189. doi:10.1146/annurev.neuro.26.041002.131052

Boudrias, M.-H., McPherson, R.L., Frost, S.B., Cheney, P.D., 2010. Output properties and organization of the forelimb representation of motor areas on the lateral aspect of the hemisphere in rhesus macaques. Cereb. Cortex 20, 169–86. doi:10.1093/cercor/bhp084

Buccino, G., 2014. Action observation treatment: a novel tool in neurorehabilitation. Philos. Trans. R. Soc. Lond. B. Biol. Sci. 369, 20130185. doi:10.1098/rstb.2013.0185

Buccino, G., Binkofski, F., Fink, G.R., Fadiga, L., Fogassi, L., Gallese, V., Seitz, R.J., Zilles, K., Rizzolatti, G., Freund, H.J., 2001. Action observation activates premotor and parietal areas in a somatotopic manner: an fMRI study. Eur. J. Neurosci. 13, 400–4.

Buccino, G., Vogt, S., Ritzl, A., Fink, G.R., Zilles, K., Freund, H.-J., Rizzolatti, G., 2004. Neural circuits underlying imitation learning of hand actions: an event-related fMRI study. Neuron 42, 323–34.

Buckner, R.L., Krienen, F.M., Castellanos, A., Diaz, J.C., Yeo, B.T.T., 2011. The organization of the human cerebellum estimated by intrinsic functional connectivity. J. Neurophysiol. 106, 2322–45. doi:10.1152/jn.00339.2011

Calmels, C., Holmes, P., Lopez, E., Naman, V., 2006. Chronometric comparison of actual and imaged complex movement patterns. J. Mot. Behav. 38, 339–48. doi:10.3200/JMBR.38.5.339-348

Carr, L., Iacoboni, M., Dubeau, M.-C., Mazziotta, J.C., Lenzi, G.L., 2003. Neural mechanisms of empathy in humans: a relay from neural systems for imitation to limbic areas. Proc. Natl. Acad. Sci. U. S. A. 100, 5497–502. doi:10.1073/pnas.0935845100

Caspers, S., Zilles, K., Laird, A.R., Eickhoff, S.B., 2010. ALE meta-analysis of action observation and imitation in the human brain. Neuroimage 50, 1148–67. doi:10.1016/j.neuroimage.2009.12.112

Catmur, C., Gillmeister, H., Bird, G., Liepelt, R., Brass, M., Heyes, C., 2008. Through the looking glass: counter-mirror activation following incompatible sensorimotor learning. Eur. J. Neurosci. 28, 1208–1215. doi:10.1111/j.1460-9568.2008.06419.x

Catmur, C., Mars, R.B., Rushworth, M.F., Heyes, C., 2011. Making Mirrors: Premotor Cortex Stimulation Enhances Mirror and Counter-mirror Motor Facilitation. J. Cogn. Neurosci. 23, 2352–2362. doi:10.1162/jocn.2010.21590

Crammond, D.J., 1997. Motor imagery: never in your wildest dream. Trends Neurosci. 20, 54–7.

Daskalakis, Z.J., Paradiso, G.O., Christensen, B.K., Fitzgerald, P.B., Gunraj, C., Chen, R., 2004. Exploring the connectivity between the cerebellum and motor cortex in humans. J. Physiol. 557, 689–700. doi:10.1113/jphysiol.2003.059808

Davare, M., Montague, K., Olivier, E., Rothwell, J.C., Lemon, R.N., 2009. Ventral premotor to primary motor cortical interactions during object-driven grasp in humans. Cortex. 45, 1050–7. doi:10.1016/j.cortex.2009.02.011

De Renzi, E., Faglioni, P., Lodesani, M., Vecchi, A., 1983. Performance of left brain-damaged patients on imitation of single movements and motor sequences. Frontal and parietal-injured patients compared. Cortex. 19, 333–43.

Debaere, F., Swinnen, S.P., Béatse, E., Sunaert, S., Van Hecke, P., Duysens, J., 2001. Brain areas involved in interlimb coordination: a distributed network. Neuroimage 14, 947–58. doi:10.1006/nimg.2001.0892

Deecke, L., Kornhuber, H.H., 1978. An electrical sign of participation of the mesial “supplementary” motor cortex in human voluntary finger movement. Brain Res. 159, 473–6.

di Pellegrino, G., Fadiga, L., Fogassi, L., Gallese, V., Rizzolatti, G., 1992. Understanding motor events: a neurophysiological study. Exp. brain Res. 91, 176–80.

Dickson, D.W., 2017. Neuropathology of Parkinson disease. Parkinsonism Relat. Disord. doi:10.1016/j.parkreldis.2017.07.033

Downing, P.E., Jiang, Y., Shuman, M., Kanwisher, N., 2001. A Cortical Area Selective for Visual Processing of the Human Body. Science (80-.). 293.

Dum, R.P., Strick, P.L., 2005. Frontal lobe inputs to the digit representations of the motor areas on the lateral surface of the hemisphere. J. Neurosci. 25, 1375–86. doi:10.1523/JNEUROSCI.3902-04.2005

Dushanova, J., Donoghue, J., 2010. Neurons in primary motor cortex engaged during action observation. Eur. J. Neurosci. 31, 386–398. doi:10.1111/j.1460-9568.2009.07067.x

Eaves, D.L., Haythornthwaite, L., Vogt, S., 2014. Motor imagery during action observation modulates automatic imitation effects in rhythmical actions. Front. Hum. Neurosci. 8, 28. doi:10.3389/fnhum.2014.00028

Eickhoff, S.B., Bzdok, D., Laird, A.R., Kurth, F., Fox, P.T., 2012. Activation likelihood estimation meta-analysis revisited. Neuroimage 59, 2349–61. doi:10.1016/j.neuroimage.2011.09.017

Eickhoff, S.B., Heim, S., Zilles, K., Amunts, K., 2006. Testing anatomically specified hypotheses in functional imaging using cytoarchitectonic maps. Neuroimage 32, 570–82. doi:10.1016/j.neuroimage.2006.04.204

Eickhoff, S.B., Laird, A.R., Grefkes, C., Wang, L.E., Zilles, K., Fox, P.T., 2009. Coordinate-based activation likelihood estimation meta-analysis of neuroimaging data: a random-effects approach based on empirical estimates of spatial uncertainty. Hum. Brain Mapp. 30, 2907–26. doi:10.1002/hbm.20718

Eickhoff, S.B., Nichols, T.E., Laird, A.R., Hoffstaedter, F., Amunts, K., Fox, P.T., Bzdok, D., Eickhoff, C.R., 2016. Behavior, sensitivity, and power of activation likelihood estimation characterized by massive empirical simulation. Neuroimage 137, 70–85. doi:10.1016/j.neuroimage.2016.04.072

Eickhoff, S.B., Paus, T., Caspers, S., Grosbras, M.-H., Evans, A.C., Zilles, K., Amunts, K., 2007. Assignment of functional activations to probabilistic cytoarchitectonic areas revisited. Neuroimage 36, 511–21. doi:10.1016/j.neuroimage.2007.03.060

Eickhoff, S.B., Stephan, K.E., Mohlberg, H., Grefkes, C., Fink, G.R., Amunts, K., Zilles, K., 2005. A new SPM toolbox for combining probabilistic cytoarchitectonic maps and functional imaging data. Neuroimage 25, 1325–35. doi:10.1016/j.neuroimage.2004.12.034

Erickson, L.C., Heeg, E., Rauschecker, J.P., Turkeltaub, P.E., 2014. An ALE meta-analysis on the audiovisual integration of speech signals. Hum. Brain Mapp. 35, 5587–605. doi:10.1002/hbm.22572

Ertelt, D., Small, S., Solodkin, A., Dettmers, C., McNamara, A., Binkofski, F., Buccino, G., 2007. Action observation has a positive impact on rehabilitation of motor deficits after stroke. Neuroimage 36 Suppl 2, T164–73. doi:10.1016/j.neuroimage.2007.03.043

Evans, A.C., Kamber, M., Collins, D.L., MacDonald, D., 1994. An MRI-Based Probabilistic Atlas of Neuroanatomy, in: Magnetic Resonance Scanning and Epilepsy. Springer US, Boston, MA pp. 263–274. doi:10.1007/978-1-4615-2546-2_48

Fadiga, L., Craighero, L., Olivier, E., 2005. Human motor cortex excitability during the perception of others’ action. Curr. Opin. Neurobiol. 15, 213–8. doi:10.1016/j.conb.2005.03.013

Ferri, S., Kolster, H., Jastorff, J., Orban, G.A., 2013. The overlap of the EBA and the MT/V5 cluster. Neuroimage 66, 412–425. doi:10.1016/j.neuroimage.2012.10.060

Fogassi, L., Ferrari, P.F., Gesierich, B., Rozzi, S., Chersi, F., Rizzolatti, G., 2005. Parietal Lobe: From Action Organization to Intention Understanding. Science (80-.). 308.

Gallese, V., Fadiga, L., Fogassi, L., Rizzolatti, G., 1996. Action recognition in the premotor cortex. Brain 119 (Pt 2), 593–609.

Grafton, S.T., Arbib, M.A., Fadiga, L., Rizzolatti, G., 1996. Localization of grasp representations in humans by positron emission tomography. 2. Observation compared with imagination. Exp. brain Res. 112, 103–11.

Grèzes, J., Decety, J., 2001. Functional anatomy of execution, mental simulation, observation, and verb generation of actions: A meta-analysis. Hum. Brain Mapp. 12, 1–19. doi:10.1002/1097-0193(200101)12:1<1::AID-HBM10>3.0.CO;2-V

Grosbras, M.-H., Paus, T., 2005. Brain Networks Involved in Viewing Angry Hands or Faces. Cereb. Cortex 16, 1087–1096. doi:10.1093/cercor/bhj050

Halsband, U., Ito, N., Tanji, J., Freund, H.J., 1993. The role of premotor cortex and the supplementary motor area in the temporal control of movement in man. Brain 116 (Pt 1.), 243–66.

Hardwick, R.M., Daglioglou, M., Miall, R.C., 2012. State estimation and the cerebellum., in: Manto, M., Gruol, D., Schmahmann, J., Koibuchi, N., Rossi, F. (Eds.), Handbook of Cerebellum and Cerebellar Disorders. Springer, Dordrecht, pp. 1297–1313.

Hardwick, R.M., Lesage, E., Eickhoff, C.R., Clos, M., Fox, P., Eickhoff, S.B., 2015. Multimodal connectivity of motor learning-related dorsal premotor cortex. Neuroimage 123, 114–28. doi:10.1016/j.neuroimage.2015.08.024

Hardwick, R.M., Rajan, V.A., Bastian, A.J., Krakauer, J.W., Celnik, P.A., 2017. Motor Learning in Stroke: Trained Patients Are Not Equal to Untrained Patients With Less Impairment. Neurorehabil. Neural Repair 31, 178–189. doi:10.1177/1545968316675432

Hardwick, R.M., Rottschy, C., Miall, R.C., Eickhoff, S.B., 2013. A quantitative meta-analysis and review of motor learning in the human brain. Neuroimage 67, 283–97. doi:10.1016/j.neuroimage.2012.11.020

Helmich, R.C., De Lange, F.P., Bloem, B.R., Toni, I., 2007. Cerebral compensation during motor imagery in Parkinson’s disease. Neuropsychologia 45, 2201–2215. doi:10.1016/j.neuropsychologia.2007.02.024

Heremans, E., Feys, P., Nieuwboer, A., Vercruysse, S., Vandenberghe, W., Sharma, N., Helsen, W., 2011. No Title. Neurorehabil. Neural Repair 25, 168–177. doi:10.1177/1545968310370750

Hétu, S., Grégoire, M., Saimpont, A., Coll, M.-P., Eugène, F., Michon, P.-E., Jackson, P.L., 2013. The neural network of motor imagery: an ALE meta-analysis. Neurosci. Biobehav. Rev. 37, 930–49. doi:10.1016/j.neubiorev.2013.03.017

Hoffstaedter, F., Grefkes, C., Caspers, S., Roski, C., Palomero-Gallagher, N., Laird, A.R., Fox, P.T., Eickhoff, S.B., 2013a. The role of anterior midcingulate cortex in cognitive motor control: Evidence from functional connectivity analyses. Hum. Brain Mapp. doi:10.1002/hbm.22363

Hoffstaedter, F., Grefkes, C., Zilles, K., Eickhoff, S.B., 2013b. The “What” and “When” of Self-Initiated Movementsd. Cereb. Cortex 23, 520–530. doi:10.1093/cercor/bhr391

Holmes, P., Calmels, C., 2008. A Neuroscientific Review of Imagery and Observation Use in Sport. J. Mot. Behav. 40, 433–445. doi:10.3200/JMBR.40.5.433-445

Ietswaart, M., Johnston, M., Dijkerman, H.C., Joice, S., Scott, C.L., MacWalter, R.S., Hamilton, S.J.C., 2011. Mental practice with motor imagery in stroke recovery: randomized controlled trial of efficacy. Brain 134, 1373–86. doi:10.1093/brain/awr077

Jackson, P.L., Lafleur, M.F., Malouin, F., Richards, C., Doyon, J., 2001. Potential role of mental practice using motor imagery in neurologic rehabilitation. Arch. Phys. Med. Rehabil. 82, 1133–1141. doi:10.1053/apmr.2001.24286

Jastorff, J., Begliomini, C., Fabbri-Destro, M., Rizzolatti, G., Orban, G.A., 2010. Coding observed motor acts: different organizational principles in the parietal and premotor cortex of humans. J. Neurophysiol. 104, 128–40. doi:10.1152/jn.00254.2010

Jeannerod, M., 2001. Neural Simulation of Action: A Unifying Mechanism for Motor Cognition. Neuroimage 14, S103–S109. doi:10.1006/nimg.2001.0832

Keysers, C., Gazzola, V., 2009. Expanding the mirror: vicarious activity for actions, emotions, and sensations. Curr. Opin. Neurobiol. 19, 666–671. doi:10.1016/j.conb.2009.10.006

Kilner, J.M., Lemon, R.N., 2013. What we know currently about mirror neurons. Curr. Biol. 23, R1057–62. doi:10.1016/j.cub.2013.10.051

Kitago, T., Krakauer, J.W., 2013. Motor learning principles for neurorehabilitation. Handb. Clin. Neurol. 110, 93–103. doi:10.1016/B978-0-444-52901-5.00008-3

Klann, J., Binkofski, F.C., Caspers, S., 2015. On the neuroanatomical parcellation and multiple functions of the inferior parietal lobule, in: Comparative Neuropsychology and Brain Imaging. pp. 75–84.

Laird, A.R., McMillan, K.M., Lancaster, J.L., Kochunov, P., Turkeltaub, P.E., Pardo, J. V, Fox, P.T., 2005. A comparison of label-based review and ALE meta-analysis in the Stroop task. Hum. Brain Mapp. 25, 6–21. doi:10.1002/hbm.20129

Lancaster, J.L., Tordesillas-Gutiérrez, D., Martinez, M., Salinas, F., Evans, A., Zilles, K., Mazziotta, J.C., Fox, P.T., 2007. Bias between MNI and Talairach coordinates analyzed using the ICBM-152 brain template. Brain Mapp. 28, 1194–1205. doi:10.1002/hbm.20345

Leek, E.C., Johnston, S.J., 2009. Functional specialization in the supplementary motor complex. Nat. Rev. Neurosci. 10, 78–78. doi:10.1038/nrn2478-c1

Lenzi, D., Trentini, C., Pantano, P., Macaluso, E., Iacoboni, M., Lenzi, G.L., Ammaniti, M., 2009. Neural basis of maternal communication and emotional expression processing during infant preverbal stage. Cereb. Cortex 19, 1124–33. doi:10.1093/cercor/bhn153

Liew, S.-L., Rana, M., Cornelsen, S., Fortunato de Barros Filho, M., Birbaumer, N., Sitaram, R., Cohen, L.G., Soekadar, S.R., 2016. Improving Motor Corticothalamic Communication After Stroke Using Real-Time fMRI Connectivity-Based Neurofeedback. Neurorehabil. Neural Repair 30, 671–675. doi:10.1177/1545968315619699

Lingnau, A., Downing, P.E., 2015. The lateral occipitotemporal cortex in action. Trends Cogn. Sci. 19, 268–277. doi:10.1016/j.tics.2015.03.006

Loporto, M., McAllister, C., Williams, J., Hardwick, R., Holmes, P., 2011. Investigating central mechanisms underlying the effects of action observation and imagery through transcranial magnetic stimulation. J. Mot. Behav. 43, 361–73. doi:10.1080/00222895.2011.604655

Lorey, B., Naumann, T., Pilgramm, S., Petermann, C., Bischoff, M., Zentgraf, K., Stark, R., Vaitl, D., Munzert, J., 2013. How equivalent are the action execution, imagery, and observation of intransitive movements? Revisiting the concept of somatotopy during action simulation. Brain Cogn. 81, 139–150. doi:10.1016/j.bandc.2012.09.011

Lotze, M., Halsband, U., 2006. Motor imagery. J. Physiol. 99, 386–395. doi:10.1016/j.jphysparis.2006.03.012

Mars, R.B., Grol, M.J., 2007. Dorsolateral Prefrontal Cortex, Working Memory, and Prospective Coding for Action. J. Neurosci. 27.

Mayka, M.A., Corcos, D.M., Leurgans, S.E., Vaillancourt, D.E., 2006. Three-dimensional locations and boundaries of motor and premotor cortices as defined by functional brain imaging: a meta-analysis1. Neuroimage 31, 1453–74. doi:10.1016/j.neuroimage.2006.02.004

Miall, R.C., 2003. Connecting mirror neurons and forward models. Neuroreport 14, 2135–7. doi:10.1097/01.wnr.0000098751.87269.77

Muckli, L., Petro, L.S., 2017. The Significance of Memory in Sensory Cortex. Trends Neurosci. 40, 255–256. doi:10.1016/j.tins.2017.03.004

Nachev, P., Kennard, C., Husain, M., 2008. Functional role of the supplementary and pre-supplementary motor areas. Nat. Rev. Neurosci. 9, 856–69. doi:10.1038/nrn2478

Nichols, T., Brett, M., Andersson, J., Wager, T., Poline, J.-B., 2005. Valid conjunction inference with the minimum statistic. Neuroimage 25, 653–60. doi:10.1016/j.neuroimage.2004.12.005

Nigel, R., Zafiris, D., Paul, F., 2015. The relationship between dorsolateral prefrontal cortical inhibition and working memory performance: a combined TMS-EEG study. Front. Hum. Neurosci. 9. doi:10.3389/conf.fnhum.2015.217.00347

Orban, G.A., 2016. Functional definitions of parietal areas in human and non-human primates. Proc. R. Soc. London B Biol. Sci. 283.

Pascual-Leone, A., Nguyet, D., Cohen, L.G., Brasil-Neto, J.P., Cammarota, A., Hallett, M., 1995. Modulation of muscle responses evoked by transcranial magnetic stimulation during the acquisition of new fine motor skills. J. Neurophysiol. 74, 1037–45.

Penfield, W., Rasmussen, T., 1950. The cerebral cortex of man. New York: Macmillan.

Picard, N., Strick, P.L., 1996. Motor areas of the medial wall: a review of their location and functional activation. Cereb. Cortex 6, 342–53.

Poldrack, R.A., 2006. Can cognitive processes be inferred from neuroimaging data? Trends Cogn. Sci. 10, 59–63. doi:10.1016/j.tics.2005.12.004

Procyk, E., Wilson, C.R.E., Stoll, F.M., Faraut, M.C.M., Petrides, M., Amiez, C., 2014. Midcingulate Motor Map and Feedback Detection: Converging Data from Humans and Monkeys. Cereb. Cortex 26, bhu213. doi:10.1093/cercor/bhu213

Rizzolatti, G., Camarda, R., Fogassi, L., Gentilucci, M., Luppino, G., Matelli, M., 1988. Functional organization of inferior area 6 in the macaque monkey. II. Area F5 and the control of distal movements. Exp. brain Res. 71, 491–507.

Rizzolatti, G., Fadiga, L., Gallese, V., Fogassi, L., 1996a. Premotor cortex and the recognition of motor actions. Brain Res. Cogn. Brain Res. 3, 131–41.

Rizzolatti, G., Fadiga, L., Matelli, M., Bettinardi, V., Paulesu, E., Perani, D., Fazio, F., 1996b. Localization of grasp representations in humans by PET: 1. Observation versus execution. Exp. brain Res. 111, 246–52.

Rottschy, C., Langner, R., Dogan, I., Reetz, K., Laird, A.R., Schulz, J.B., Fox, P.T., Eickhoff, S.B., 2012. Modelling neural correlates of working memory: a coordinate-based meta-analysis. Neuroimage 60, 830–46. doi:10.1016/j.neuroimage.2011.11.050

Rushworth, M.F., Nixon, P.D., Wade, D.T., Renowden, S., Passingham, R.E., 1998. The left hemisphere and the selection of learned actions. Neuropsychologia 36, 11–24.

Schlerf, J.E., Verstynen, T.D., Ivry, R.B., Spencer, R.M.C., 2010. Evidence of a Novel Somatopic Map in the Human Neocerebellum During Complex Actions. J. Neurophysiol. 103.

Shadmehr, R., Krakauer, J.W., 2008. A computational neuroanatomy for motor control. Exp. Brain Res. 185, 359–381. doi:10.1007/s00221-008-1280-5

Sirigu, A., Duhamel, J.-R., Cohen, L., Pillon, B., Dubois, B., Agid, Y., 1996. The Mental Representation of Hand Movements After Parietal Cortex Damage. Science (80-.). 273.

Sommer, M.A., 2003. The role of the thalamus in motor control. Curr. Opin. Neurobiol. 13, 663–70.

Stippich, C., Ochmann, H., Sartor, K., 2002. Somatotopic mapping of the human primary sensorimotor cortex during motor imagery and motor execution by functional magnetic resonance imaging. Neurosci. Lett. 331, 50–54.

Tanaka, S., Honda, M., Sadato, N., 2005. Modality-specific cognitive function of medial and lateral human Brodmann area 6. J. Neurosci. 25, 496–501. doi:10.1523/JNEUROSCI.4324-04.2005

Tkach, D., Reimer, J., Hatsopoulos, N.G., 2007. Congruent Activity during Action and Action Observation in Motor Cortex. J. Neurosci. 27.

Turkeltaub, P.E., Eden, G.F., Jones, K.M., Zeffiro, T.A., 2002. Meta-analysis of the functional neuroanatomy of single-word reading: method and validation. Neuroimage 16, 765–80.

Turkeltaub, P.E., Eickhoff, S.B., Laird, A.R., Fox, M., Wiener, M., Fox, P., 2012. Minimizing within-experiment and within-group effects in Activation Likelihood Estimation meta-analyses. Hum. Brain Mapp. 33, 1–13. doi:10.1002/hbm.21186

Turner, R.S., Desmurget, M., Grethe, J., Crutcher, M.D., Grafton, S.T., 2003. Motor subcircuits mediating the control of movement extent and speed. J. Neurophysiol. 90, 3958–66. doi:10.1152/jn.00323.2003

Urgesi, C., Candidi, M., Ionta, S., Aglioti, S.M., 2007. Representation of body identity and body actions in extrastriate body area and ventral premotor cortex. Nat. Neurosci. 10, 30–31. doi:10.1038/nn1815

van der Gaag, C., Minderaa, R.B., Keysers, C., 2007. Facial expressions: what the mirror neuron system can and cannot tell us. Soc. Neurosci. 2, 179–222. doi:10.1080/17470910701376878

Vigneswaran, G., Philipp, R., Lemon, R.N., Kraskov, A., 2013. M1 Corticospinal Mirror Neurons and Their Role in Movement Suppression during Action Observation. Curr. Biol. 23, 236–243. doi:10.1016/j.cub.2012.12.006

Vogt, S., Di Rienzo, F., Collet, C., Collins, A., Guillot, A., 2013. Multiple roles of motor imagery during action observation. Front. Hum. Neurosci. 7, 807. doi:10.3389/fnhum.2013.00807

Voon, V., Baek, K., Enander, J., Worbe, Y., Morris, L.S., Harrison, N.A., Robbins, T.W., Rück, C., Daw, N., 2015. Motivation and value influences in the relative balance of goal-directed and habitual behaviours in obsessive-compulsive disorder. Transl. Psychiatry 5, e670. doi:10.1038/tp.2015.165

Ward, J., Banissy, M.J., 2015. Explaining mirror-touch synesthesia. Cogn. Neurosci. 6, 118–133. doi:10.1080/17588928.2015.1042444

Weiss, P.H., Rahbari, N.N., Hesse, M.D., Fink, G.R., 2008. Deficient sequencing of pantomimes in apraxia. Neurology 70, 834–40. doi:10.1212/01.wnl.0000297513.78593.dc

Williams, S.E., Guillot, A., Di Rienzo, F., Cumming, J., 2015. Comparing self-report and mental chronometry measures of motor imagery ability. Eur. J. Sport Sci. 15, 703–11. doi:10.1080/17461391.2015.1051133

Wise, S.P., Boussaoud, D., Johnson, P.B., Caminiti, R., 1997. Premotor and parietal cortex: corticocortical connectivity and combinatorial computations. Annu. Rev. Neurosci. 20, 25–42. doi:10.1146/annurev.neuro.20.1.25

Wise, S.P., Murray, E.A., 2000. Arbitrary associations between antecedents and actions. Trends Neurosci. 23, 271–6.

Wollenweber, F.A., Halfter, S., Brügmann, E., Weinberg, C., Cieslik, E.C., Müller, V.I., Hardwick, R.M., Eickhoff, S.B., 2014. Subtle cognitive deficits in severe alcohol addicts - Do they show a specific profile? J. Neuropsychol. 8, 147–153. doi:10.1111/jnp.12001

Xu, J., Ejaz, N., Hertler, B., Branscheidt, M., Widmer, M., Faria, A. V., Harran, M.D., Cortes, J.C., Kim, N., Celnik, P.A., Kitago, T., Luft, A.R., Krakauer, J.W., Diedrichsen, J., 2017. Separable systems for recovery of finger strength and control after stroke. J. Neurophysiol. 118.

Yousry, T.A., Schmid, U.D., Alkadhi, H., Schmidt, D., Peraud, A., Buettner, A., Winkler, P., 1997. Localization of the motor hand area to a knob on the precentral gyrus. A new landmark. Brain 120 (Pt 1), 141–57.

Zabicki, A., de Haas, B., Zentgraf, K., Stark, R., Munzert, J., Krüger, B., 2017. Imagined and Executed Actions in the Human Motor System: Testing Neural Similarity Between Execution and Imagery of Actions with a Multivariate Approach. Cereb. Cortex 27, 4523–4536. doi:10.1093/cercor/bhw257

Zeiler, S.R., Hubbard, R., Gibson, E.M., Zheng, T., Ng, K., O’Brien, R., Krakauer, J.W., 2015. Paradoxical Motor Recovery From a First Stroke After Induction of a Second Stroke: Reopening a Postischemic Sensitive Period. Neurorehabil. Neural Repair. doi:10.1177/1545968315624783

Zeiler, S.R., Krakauer, J.W., 2013. The interaction between training and plasticity in the poststroke brain. Curr. Opin. Neurol. 26, 609–16. doi:10.1097/WCO.0000000000000025

Zhang, X., de Beukelaar, T.T., Possel, J., Olaerts, M., Swinnen, S.P., Woolley, D.G., Wenderoth, N., 2011. Movement observation improves early consolidation of motor memory. J. Neurosci. 31, 11515–20. doi:10.1523/JNEUROSCI.6759-10.2011

